# An Nrf2-Nup214 positive feedback loop sustains the antioxidant response and promotes microbial infection in ticks

**DOI:** 10.64898/2026.05.06.723311

**Authors:** Kaylee A. Vosbigian, Chelsea A. Osbron, Brianna P. Steiert, Kristin L. Rosche, Sarah J. Wright, Elisabeth Ramirez-Zepp, Dana K. Shaw

## Abstract

*Ixodes scapularis* ticks are obligate hematophagous ectoparasites that take a single bloodmeal per life stage. During blood digestion, large quantities of heme and iron are released, leading to the production of reactive oxygen species. To counter the oxidative stress, ticks have evolved robust antioxidant systems. We previously found that the antioxidant transcription factor Nrf2 is activated during infection and supports *Anaplasma phagocytophilum* colonization in ticks. To investigate the Nrf2 regulatory network, we queried promoter regions in the *Ixodes* genome for Nrf2 binding sites. The gene encoding the nuclear pore complex protein *nucleoporin 214* was identified as a target that is induced during *Anaplasma* infection and putatively regulated by Nrf2. *Ixodes nup214* was experimentally validated as an Nrf2-regulated gene through functional luciferase reporter assays, pharmacological manipulation, and RNA interference transcriptional repression. We found that elevated expression of Nup214 leads to binding of the nuclear export protein CRM1, thereby trapping Nrf2 in the nucleus. Increased nuclear retention of Nrf2 prolonged the antioxidant response and functionally supported *Anaplasma* survival. Our findings uncover a previously unknown mechanism potentiated by Nrf2 that supports *Anaplasma* infection in ticks.

**SIGNIFICANCE:** Ticks have evolved to rely on host blood for nutrition and development. However, blood feeding comes with drawbacks. During digestion, large quantities of heme and iron are released, which promote the production of reactive oxygen species. To counter this, ticks have evolved robust antioxidant responses to protect themselves against oxidative stress. How tick-transmitted microbes, such as *Anaplasma phagocytophilum*, withstand the oxidative challenge in the tick is not well understood. Here, we demonstrate that, during infection, host-driven antioxidant defenses coordinated by the transcription factor Nrf2 support *Anaplasma* infection and persistence in ticks. We show that Nrf2 upregulates the nuclear pore complex protein Nucleoporin 214, which sequesters Nrf2 in the nucleus. The prolonged nuclear retention of Nrf2 enhances the tick antioxidant response, thereby supporting *Anaplasma* survival in the arthropod vector.

## INTRODUCTION

In the United States, the incidence of tick-borne disease is rising as ticks expand their ecological ranges^1–3^. *Ixodes scapularis,* the North American deer tick, can transmit at least seven pathogens that affect human health, including *Anaplasma phagocytophilum, Borrelia burgdorferi, Borrelia mayonii, Borrelia miyamotoi, Babesia microti, Ehrlichia muris eauclariensis,* and *Powassan virus*^4^. These pathogens cause stress to the arthropod vector by hijacking nutrients and producing toxic byproducts^5–7^. Additionally, the parasitic lifestyle of ticks comes with unique physiological stressors. As obligate hematophagous arthropods, *I. scapularis* ticks depend on host blood for survival; however, rapid temperature shifts and high concentrations of dietary heme and iron impart molecular stress^8–12^. To mitigate these physiological strains, cellular stress responses are activated that restore homeostasis^5,13–24^.

We previously demonstrated that the unfolded protein response (UPR) impacts colonization and persistence of the obligate intracellular pathogen *A. phagocytophilum* in *I. scapularis* ticks^7,13,22^. The UPR is a conserved cellular stress response network consisting of three endoplasmic reticulum (ER) stress sensors: PERK, ATF6, and IRE1α^24–26^. Accumulation of unfolded proteins in the ER activates the UPR, leading to downstream signaling cascades that are generally involved in restoring cellular homeostasis^24^. Recently, we found that the PERK branch of the UPR supports *A. phagocytophilum* colonization in ticks by activating Nrf2, a member of the Cap’n’Collar family of basic leucine zipper (bZIP) transcription factors^13,27,28^. This potentiates an antioxidant response that benefits the bacterium^13^. However, the *Ixodes* genome notably lacks the canonical Nrf2-regulated genes *heme oxygenase 1* (*HMOX1*) and *NAD(P)H quinone dehydrogenase 1* (*NQO1*), which are central to the antioxidant response in mammals^29,30^. This suggests that the Nrf2-regulated gene network in ticks is unique and diverges from what has been characterized in model organisms. The genes regulated by Nrf2, and how this supports *Anaplasma* colonization and persistence in the arthropod, remain unknown.

Nrf2 drives transcription by binding to DNA sequence motifs termed Antioxidant Response Elements (AREs), which are found in the promoter region of targeted genes^31,32^. Herein, we predict the Nrf2 regulatory network in *I. scapularis* ticks by querying promoter regions for Nrf2 binding motifs. *Nucleoporin 214* (*nup214*) was identified and validated as an Nrf2-regulated gene that supports *A. phagocytophilum* colonization in *I. scapularis.* Through pharmacological manipulations, RNA interference (RNAi), and cellular fractionation, we demonstrate that Nup214 enhances nuclear accumulation of Nrf2 during infection. This functionally enhances the antioxidant response and supports *A. phagocytophilum* survival by neutralizing reactive oxygen species (ROS). Collectively, we show a previously unknown positive feedback loop between Nrf2 and Nup214 that functionally enhances antioxidant responses and promotes microbial infection in the arthropod.

## RESULTS

### Identifying putative Nrf2-regulated genes in ticks

Previously, we found that *Ixodes* Nrf2 is activated by *B. burgdorferi* and *A. phagocytophilum* infection and supports bacterial colonization^13^. Under homeostatic conditions, the negative regulator Keap1 (Kelch ECH associating protein 1) forms a complex with Nrf2 in the cytoplasm, which targets it for degradation^30,33,34^. Under activating conditions, Nrf2 is phosphorylated by PERK^35^, which causes Keap1 to disassociate. Alternatively, redox-sensing residues on Keap1 can be directly modified by ROS, also causing it to disassociate from Nrf2^33,36^. Free Nrf2 then translocates to the nucleus and functions as a transcription factor (Fig 1A)^13,30,35,37^. To determine the subcellular localization of *Ixodes* Nrf2 during *Anaplasma* infection, we probed nuclear fractions with an Nrf2-specific antibody and found that infected ISE6 tick cells showed increased nuclear accumulation (Fig 1B, Fig S1A-B). To determine if increased Nrf2 nuclear accumulation results in functional gene expression differences, we quantified the expression of the Nrf2-regulated gene, *glutathione s-transferase (gst) theta-1*^38^. We found that *Anaplasma-*infected tick cells exhibited higher levels of *gst theta-1* expression when compared to uninfected cells (Fig 1C, Fig S2), indicating that *Ixodes* Nrf2 is activated and functionally drives a transcriptional response during *Anaplasma* infection.

**Figure 1.**
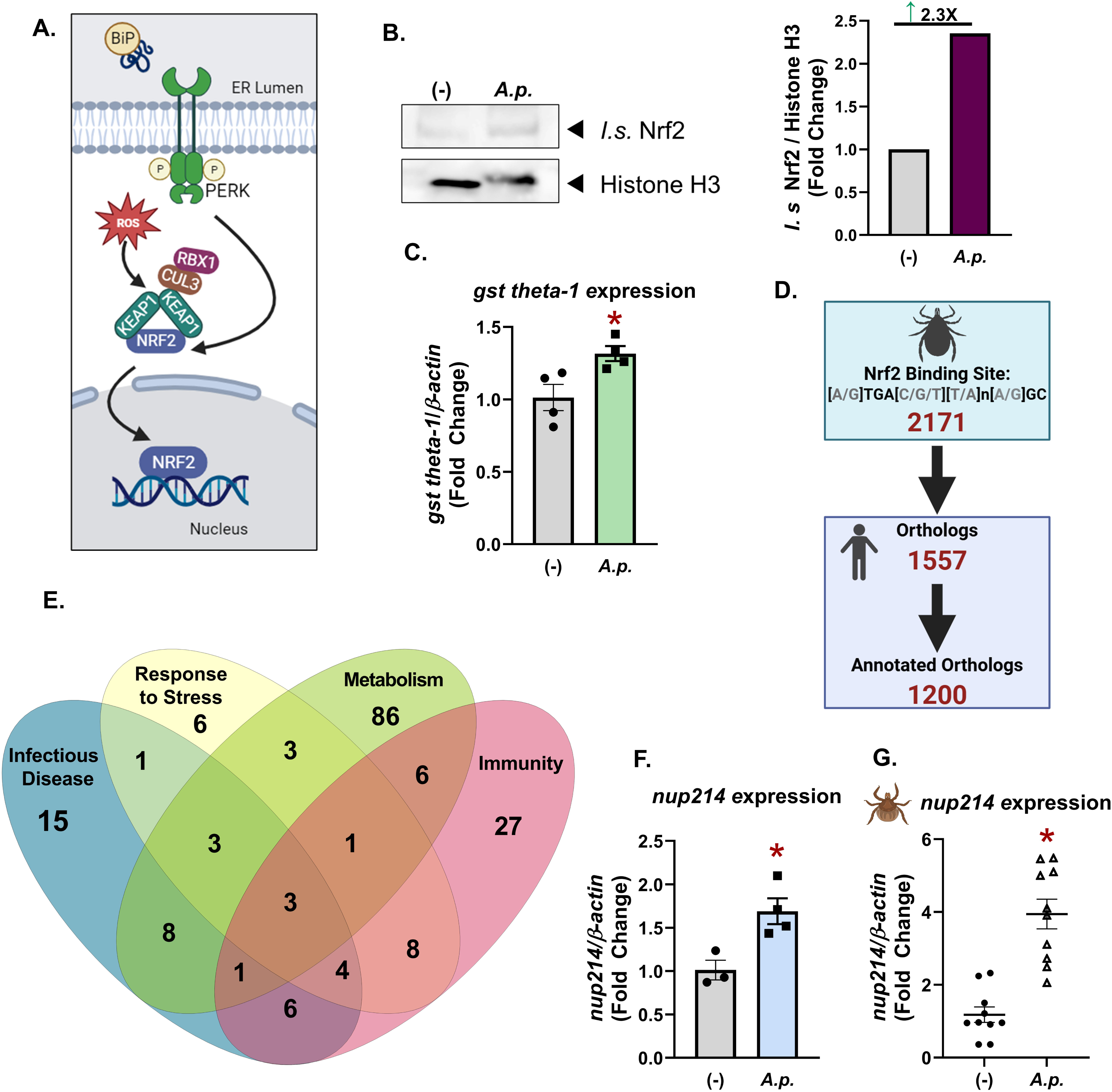
The predicted Nrf2-regulated network in *I. scapularis*. (A) Schematic depicting Nrf2 activation. (B) Nrf2 (∼150 kDa) immunoblot against nuclear extracts of ISE6 cells infected with *A. phagocytophilum* (MOI 50) for 6 hours. Histone H3 (∼16 kDa) was used as a loading control. Protein expression differences were quantified by ImageJ and are expressed as a ratio of Nrf2 to Histone H3. (C) Expression of *gst theta-1* in ISE6 cells infected with *A. phagocytophilum* compared to uninfected. (D) Schematic of results from Nrf2 binding search. (E) Reactome pathways represented in putative Nrf2-regulated genes in *I. scapularis* genome. Expression of *nucleoporin 214* from uninfected and *Anaplasma-* infected (F) ISE6 cells and (G) *I. scapularis* larvae rested for 7 days post-repletion. *, *P* < 0.05 (Student’s *t* test).

Nrf2 is a master regulator of the antioxidant response and is associated with the direct transcriptional regulation of antioxidant genes^29,30,39,40^. In addition to the antioxidant response, Nrf2 also regulates transcriptional networks orchestrating many cellular processes including metabolism, inflammation, autophagy, proteostasis, mitochondrial biogenesis, and immune signaling^30,39,41^. Canonical Nrf2-regulated genes include *GSTs, heme oxygenase 1* (*HMOX1*), and *NAD(P)H quinone dehydrogenase 1* (*NQO1*), which are central to the antioxidant responses in mammals^29,30,40–42^. However, the *Ixodes* genome notably lacks *HMOX1* and *NQO1*^43–45^, suggesting that the Nrf2 regulatory network in ticks is unique and diverges from what has been characterized in mammals.

Previously we found that human and *Ixodes* Nrf2s are structurally similar and that amino acids mediating DNA interactions within the basic leucine zipper (bZIP) domain are well conserved^13^. This conservation provided the rationale for interrogating the *Ixodes* genome for Nrf2-regulated genes. We used our previously reported web-based tool, Arthroquest^7^, to scan promoter regions of the *I. scapularis* genome for Nrf2 binding sites [A/G]TGA[C/G/T][T/A]n[A/G]GC (where “n” indicates any nucleotide)^37^. We found that, out of 34,235 genes in the *I. scapularis* genome, 2,171 genes are putatively regulated by Nrf2 (Fig 1D, Supplementary Dataset 1).

We next sought to analyze the *Ixodes* Nrf2-regulated gene network. Relative to model organisms, there is limited gene information associated with the *I. scapularis* genome. To circumvent this limitation, we identified human orthologs for the 2,171 tick genes. A total of 1,557 tick genes showed significant orthology to human genes and 1,200 had associated Gene Ontology terms or Reactome annotations (Fig 1D). Of these, 415 orthologs were classified as being involved in Nrf2-regulated pathways (Fig S3, Supplementary Dataset 1). Functional classification of these orthologs revealed associations with metabolism (111; R-HSA-1430728), the immune system (56; R-HSA-168256), infectious diseases (41; R-HSA-5663205), and response to stress (29; R-HSA-2262752) (Fig 1E). Three proteins were predicted to be involved in all four pathways: the 26S proteosome non-ATPase regulatory subunit 13 (PSMD13) and two nuclear pore proteins, Nucleoporin 214 (Nup214) and Nucleoporin TPR (Translocated Promoter Region). PSMD13 is a subunit of the 26S proteosome involved in non-lysosomal protein degradation of misfolded or damaged proteins^46^. Nup214 is a phenylalanine-glycine (FG) repeat protein anchored to the cytoplasmic face of the nuclear pore complex (NPC) and is involved in regulating nuclear transport^47–50^. Nucleoporin TPR is located in the nuclear basket of the pore complex and is also involved in regulating nuclear transport^48,51^.

To prioritize putative Nrf2-regulated gene candidates, we surveyed published transcriptomic data sets from *Anaplasma-*infected *I. scapularis* midguts^52^. Out of the three candidates, we found that only *nup214* was significantly increased in infected ticks^52^. We experimentally confirmed this by quantifying *nup214* expression in *A. phagocytophilum-*infected ISE6 tick cells (Fig 1F) and whole *I. scapularis* larvae (Fig 1G). Nrf2 has never been linked to regulating the NPC, but it does require nuclear entry to orchestrate transcriptional processes. Given that Nup214 is specifically targeted by some viral^53–55^ and bacterial^56^ pathogens to facilitate infection, we hypothesized that this may be another mechanism by which Nrf2 is supporting *Anaplasma*.

### Tick Nucleoporin 214 is regulated by Nrf2 during infection

Once in the nucleus, Nrf2 binds to AREs in a heterodimer with MafG (small musculo-aponeurotic fibrosarcoma protein G)^57^. To predict structural interactions between *I. scapularis* Nrf2, MafG, and the *nup214* promoter, we used AlphaFold3^58^ (Fig 2A; Fig S4). While in a heterodimer with MafG, the prediction shows direct interaction between the DNA binding residues of Nrf2^13^ (green; R877, R880, R882, N885, A888, A889,R893, R895, and K896) and the Nrf2 DNA binding motif (orange) found in the *I. scapularis nup214* promoter (Fig 2A). We next evaluated if *nup214* transcription correlated with *nrf2* expression during *A. phagocytophilum* infection. We used RNA interference (RNAi) to transcriptionally silence *nrf2* in tick cells and found that, under these conditions, *nup214* expression was also significantly decreased (Fig 2B). Conversely, silencing the negative regulator of Nrf2, *keap1,* led to increased *nup214* expression (Fig 2C), suggesting that Nrf2 positively regulates *nup214*.

**Figure 2.**
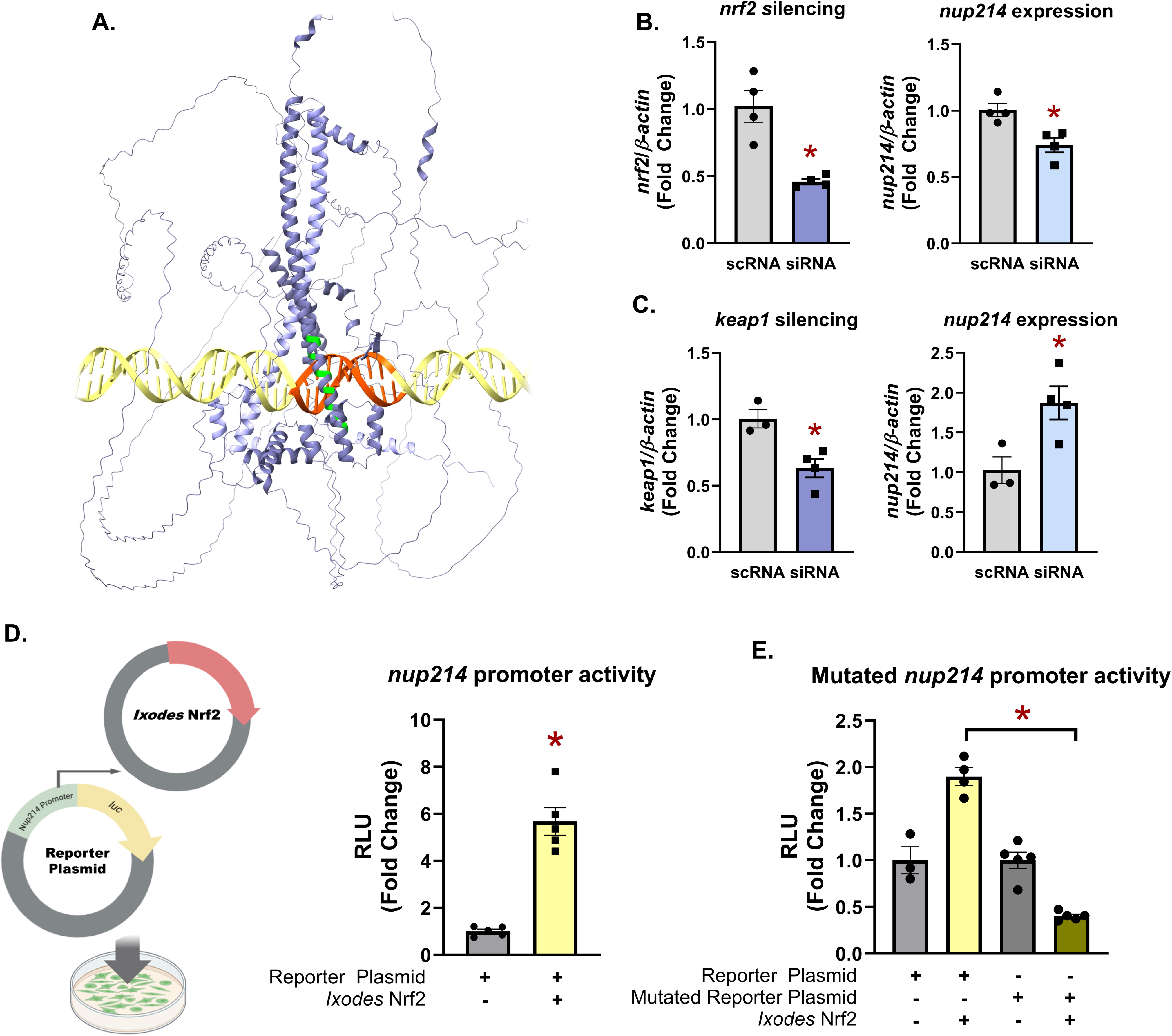
*nucleoporin 214* expression is regulated by Nrf2. (A) The AlphaFold3 predicted DNA-protein complex with *I. scapularis* Nrf2 (dark purple), *I. scapularis* MafG (light purple), and the predicted promoter of *nup214* (yellow). DNA binding residues are highlighted in green and the consensus binding site is indicated in orange. (B) *nup214* expression quantified from *A. phagocytophilum* infected IDE12 tick cells treated with siRNA targeting *nrf2* or a scrambled control. (C) *nup214* expression quantified from *A. phagocytophilum* infected ISE6 cells treated with siRNA targeting *keap1* or a scrambled control. (D-E) HEK 293T cells were co-transfected with plasmids constitutively expressing *I. scapularis* Nrf2 and (D) a luciferase reporter plasmid with the *I. scapularis nup214* promoter or (E) a luciferase reporter plasmid with a mutated *I. scapularis nup214* promoter. Luciferase activity was normalized to the condition with only the luciferase reporter plasmid transfected. RLU is relative luminescence units. *, *P* < 0.05 (Student’s *t* test). scRNA, scrambled RNA; siRNA, small interfering RNA.

To experimentally validate that Nrf2 drives expression from the *nup214* promoter, we designed a custom luciferase reporter assay. A reporter plasmid was engineered with the *Ixodes nup214* promoter upstream from a gene encoding *luciferase*. We next constructed a plasmid that constitutively expressed *Ixodes* Nrf2. Both plasmids were co-transfected into HEK 293T cells for 24 hours (Fig 2D). A positive control plasmid containing *luciferase* driven by a known *nrf2* promoter was also co-transfected with the *Ixodes* Nrf2 expression construct (Fig S5). After 24 hours, D-luciferin was added and luciferase activity was quantified. We found that luciferase activity significantly increased when the cells were expressing *I. scapularis* Nrf2, demonstrating *nup214* promoter activity (Fig 2D). To confirm this was specifically dependent on *Ixodes* Nrf2, we generated a reporter plasmid containing the *nup214* promoter with a mutated Nrf2 binding site (GTGATTAAGC to TGCTAGCTTA) (Fig 2E). When the mutated *nup214* promoter plasmid was co-transfected with *Ixodes* Nrf2, luciferase activity was completely abolished, indicating that Nrf2 specifically binds the *nup214* promoter to drive transcription (Fig 2E). Overall, these data demonstrate that *I. scapularis* Nrf2 positively regulates the expression of *nup214*.

### Anaplasma infection is supported by Ixodes nup214

Given that *Ixodes nup214* is regulated by Nrf2 and that its expression increases during *Anaplasma* infection (Fig 1F-G), we next asked how *nup214* impacts bacterial colonization in ticks. To address this, we silenced *nup214* in ISE6 tick cells using RNAi and then infected with *A. phagocytophilum* for 24 hours. We found that blocking *nup214* expression led to decreased *A. phagocytophilum* (Fig 3A), similar to the phenotype observed when *nrf2* is knocked down^13^. To determine if a similar phenotype was observed *in vivo*, we silenced *nup214* expression by immersing *I. scapularis* larvae in silencing RNA or in non-specific scrambled controls. The larvae then fed to repletion on *A. phagocytophilum-*infected mice. Pathogen numbers in the ticks were quantified at two different time points that correspond to (i) pathogen acquisition (immediately after repletion) or (ii) population expansion in the tick (seven days post-repletion). We found that decreasing the expression of *nup214* did not have an impact on *A. phagocytophilum* acquisition (Fig 3B-C). However, silencing *nup214* was detrimental to pathogen population expansion in the tick after seven days (Fig 3D), suggesting that *nup214* supports *A. phagocytophilum* colonization, growth, and dissemination in ticks.

**Figure 3.**
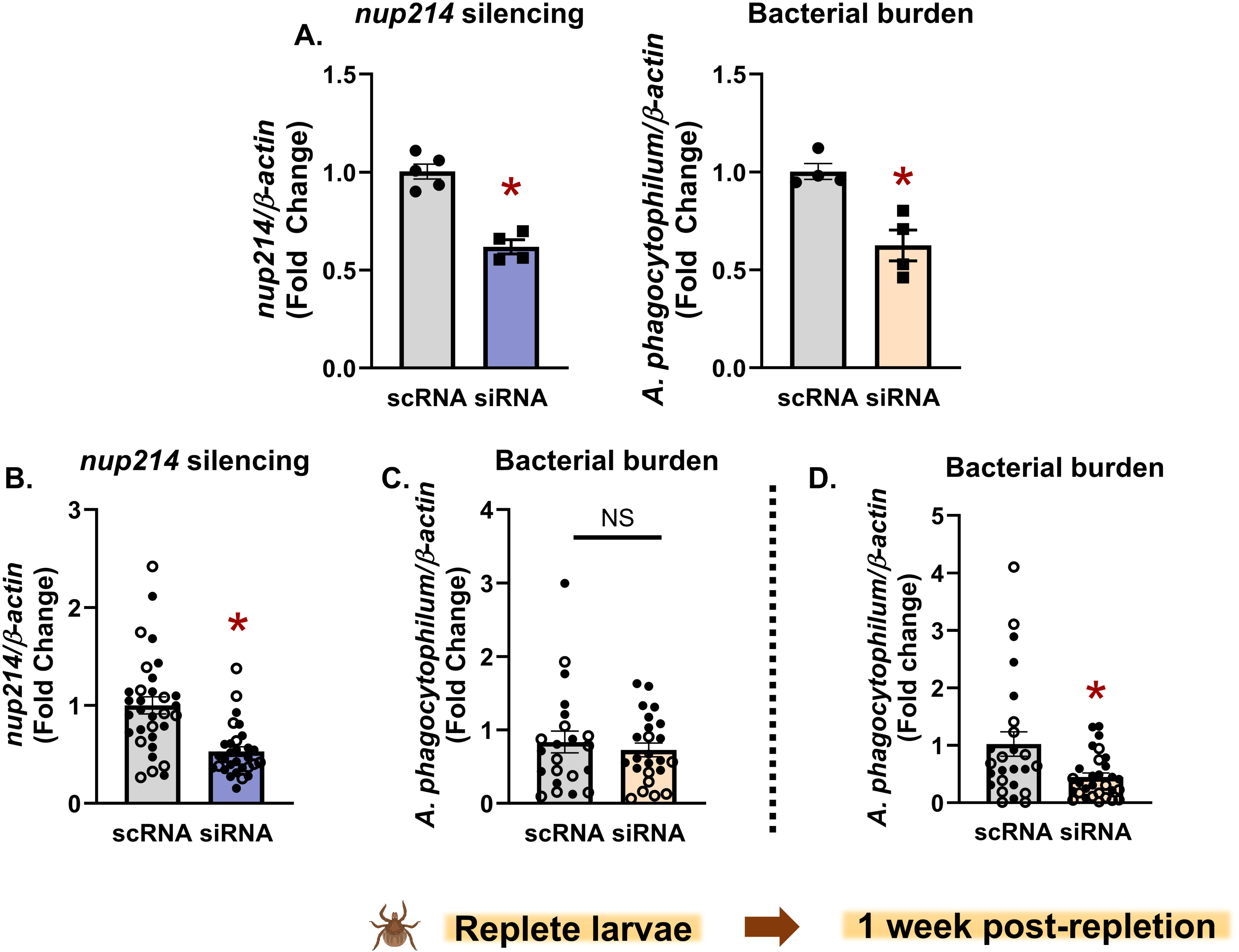
*Anaplasma* infection and colonization is supported by *Ixodes* Nup214. (A) Tick cells were transfected with siRNA or scrambled control (scRNA) prior to infecting with *A. phagocytophilum* (MOI 50) for 18-24 hours. (B) Larvae were treated with either siRNA for *nup214* or scRNA before being fed on *A. phagocytophilum*-infected mice. Larvae were either (C) immediately processed after feeding or (D) rested for 7 days. Each data point is representative of 1 larva. Open and closed dots represent experimental replicates. Silencing efficiency and bacterial burden were measured by qRT-PCR. *, *P* < 0.05 (Student’s *t* test). scRNA, scrambled RNA; siRNA, small interfering RNA. NS, not significant.

### Nup214-CRM1-mediated nuclear accumulation of Nrf2 supports Anaplasma

Nup214 has been extensively studied in model systems. It functions as a phenylalanine-glycine (FG) repeat nucleoporin that is involved in nuclear import and export of proteins^50,59–62^. Although *Ixodes* and mammalian Nup214 share only 33% sequence identity (Fig S6A), the N-terminal beta-propeller domain, which is responsible for binding to the nuclear pore complex (NPC)^63,64^, is structurally conserved in the AlphaFold3 prediction (Fig S6B-C). This suggests that *Ixodes* Nup214 would be similarly positioned on the nucleus. Importantly, *Ixodes* Nup214 also contains the unstructured FG repeat region that is involved in regulating nuclear transport, suggesting a conserved function with human Nup214 (Fig S6A, highlighted in green).

Nup214 regulates protein export through interactions with the nuclear transport receptor CRM1 (Chromosome Region Maintenance 1; also known as Exportin 1)^47,49,59,65,66^. CRM1 recognizes nuclear export signals (NESs) and facilitates the transfer of proteins across the nuclear membrane^67,68^. If Nup214 expression is high enough, it can sequester CRM1 on the cytoplasmic side of the nucleus, thereby blocking CMR1-mediated export of transcription factors^49,66^. Mammalian Nrf2 is dependent on CRM1-mediated export and, when this is pharmacologically repressed, Nrf2 accumulates in the nucleus^69,70^. For this reason, we asked whether *Ixodes* Nup214 expression affects Nrf2 nuclear accumulation through CRM1. We first evaluated if *Ixodes* Nrf2 and CRM1 are predicted to interact. The *Ixodes* Nrf2 sequence was scanned for predicted NESs using the LocNES program^71^ and several were identified (Fig S7A). We also found significant conservation between *Ixodes* and human CRM1 (75.42% identity, Fig S7B-D) and, when modeled in AlphaFold3, *Ixodes* CRM1 and the NES in the bZIP domain of *Ixodes* Nrf2 are predicted to interact (Fig S8).

To experimentally validate that *Ixodes* CRM1 and Nrf2 bind, we performed reciprocal pulldowns. Owing to the significant size of both Nrf2 (107 kDa) and CRM1 (126 kDa), we chose to express truncated forms of the proteins with the portions that are predicted to interact. We used a FLAG-tagged, C-terminal portion of *Ixodes* CRM1 (aa1603-3310) that contains the NES binding groove^72^ (Fig S7B) and an HA-tagged, C-terminal portion of Nrf2 (aa 2101-3003) that contains the conserved NES (Fig S7A) (Fig 4A). Using these constructs, HEK 293T cells were either singly or co-transfected. Protein lysates were immunoprecipitated with FLAG antibody conjugated agarose, which showed that CRM1 pulls down Nrf2 (Fig 4A). The reciprocal approach with antibodies against the HA tag revealed the same result, demonstrating that *Ixodes* CRM1 and Nrf2 specifically interact.

**Figure 4.**
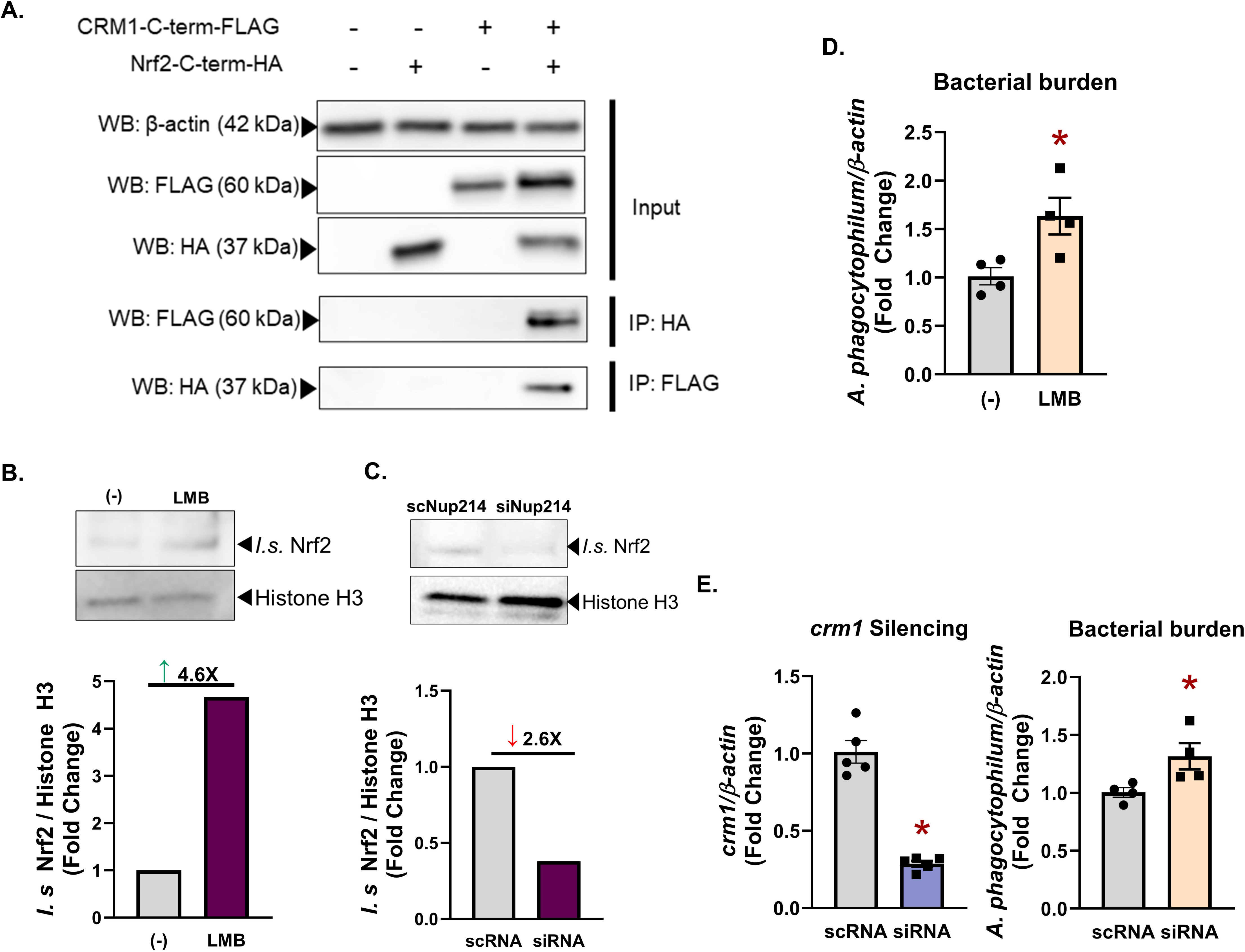
CRM1 impacts Nrf2 nuclear accumulation and *Anaplasma* infection in ticks. (A) Western blot of immunoprecipitation (IP) analysis showing interaction between HA-tagged *Ixodes* Nrf2 (aa 701-1000) and FLAG-tagged *Ixodes* CRM1 (aa 1603-3310) expressed in HEK 293T cells. Nuclear extracts of tick cells treated with (B) 20nM LMB compared to the no treatment control (-) or (C) siRNA for *nup214* compared to the scrambled control. Immunoblots probed for Nrf2 (∼150 kDa) and Histone H3 (16 kDa). Protein expression differences were quantified by ImageJ and are expressed as a ratio of Nrf2 to Histone H3. (D) Tick cells infected with *A. phagocytophilum* (MOI 50) for 18 hours and treated with LMB (20 nM) for 3 hours compared to control (-). (E) Tick cells were transfected with siRNA for *crm1* or scrambled control (scRNA) prior to infecting with *A. phagocytophilum* (MOI 50) for 18-24 hours. Silencing and bacterial burdens were quantified by qRT-PCR. Data are representative of at least three to five biological replicates and two technical replicates. Error bars show SEM. *, *P* < 0.05 (Student’s *t* test). scRNA, scrambled RNA; siRNA, small interfering RNA.

To examine the functional impact of CRM1 on Nrf2 localization, we treated ISE6 tick cells with the CRM1 inhibitor Leptomycin B (LMB), which allosterically blocks binding to NES-containing proteins^73^. We first confirmed that LMB does not affect tick cell viability (Fig S9). We then isolated nuclear fractions from LMB-treated tick cells and analyzed them by immunoblot with an antibody specific for Nrf2. We found that there was more nuclear accumulation of Nrf2 when compared to untreated controls (Fig 4B). Since Nup214 can interfere with CRM1’s function by sequestering it on the cytoplasmic side of the nucleus^49,66^, we next asked if knocking down *nup214* in tick cells would inhibit nuclear accumulation of Nrf2. To address this, we silenced *nup214* for 48 hours and then isolated the nuclei. Immunoblotted nuclear fractions showed decreased Nrf2 accumulation when *nup214* is silenced (Fig 4C). This suggests that when Nup214 is not present to bind and sequester CRM1, NES-containing proteins can freely be exported from the nucleus.

Since Nrf2 supports *Anaplasma* colonization and growth in ticks, we next asked if CRM1 function and expression impacts bacterial numbers. Tick cells were treated with LMB and infected with *Anaplasma* for 24 hours. We found that inhibiting CRM1 through LMB treatment caused a significant increase in bacterial numbers (Fig 4D). Similarly, when tick cells were treated with silencing RNA targeting *crm1,* there was a corresponding increase in *Anaplasma* compared to the scrambled control (Fig 4E). These data collectively demonstrate that inhibiting CRM1-mediated nuclear export causes nuclear accumulation of Nrf2 and is beneficial to *A. phagocytophilum*.

### Nup214 potentiates the antioxidant response in ticks

Given that Nrf2 orchestrates the expression of genes involved in antioxidant defenses, we asked if Nup214-dependent nuclear accumulation of Nrf2 impacts the oxidative state of tick cells during infection. Cells were treated with silencing RNAs targeting *nup214* and infected with *A. phagocytophilum* for 24 hours. ROS were then quantified. In alignment with our previous report^13^, we observed that *A. phagocytophilum* infection increased ROS in tick cells (Fig 5A). When *nup214* was silenced, we found that ROS accumulation in infected tick cells was exacerbated (Fig 5A). To determine if this was attributable to decreased Nrf2 transcriptional activity, we measured expression of the Nrf2-regulated antioxidant gene *gst theta-1*^41,74^. We found that knocking down *nup214* during *Anaplasma* infection led to a decrease in *gst theta-1* expression (Fig 5B). We next assayed the enzymatic antioxidant activity of GST. We found that *Anaplasma* infection increased antioxidant activity (Fig 5C), as expected. However, when *nup214* was silenced, GST activity was completely abolished (Fig 5C). These data demonstrate that Nup214 amplifies the antioxidant response during *Anaplasma* infection through Nrf2 nuclear accumulation.

**Figure 5.**
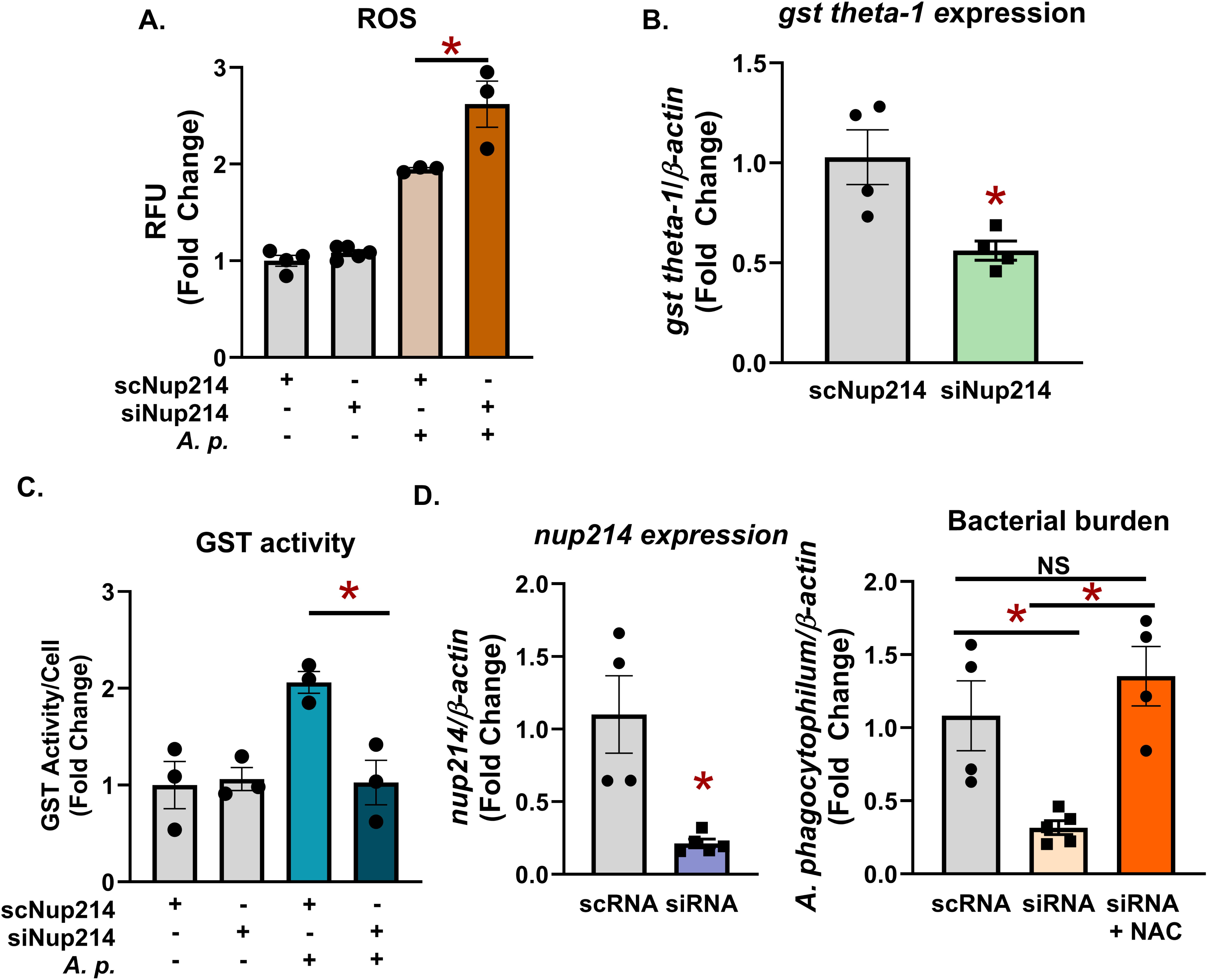
Nup214 expression amplifies the antioxidant response. (A) ROS measurement in ISE6 cells infected and treated with siRNA for *nup214* compared to uninfected, scrambled control. (B) *gst theta-1* expression and (C) GST activity in ticks cells treated with siRNA for *nup214* during infection compared to the scrambled control. (D) ISE6 tick cells with *nup214* silenced, infected with *Anaplasma,* and in the presence of NAC or alone for 24 hours. Silencing and bacterial burdens were quantified by qRT-PCR. Data are representative of at least three to five biological replicates and two technical replicates. Error bars show SEM. *, *P <* 0.05 (Student’s *t* test). NAC, N-acetyl cysteine. RFU, relative fluorescence unit. scRNA, scrambled RNA; siRNA, small interfering RNA.

It is well established that *Anaplasma* is susceptible to the bactericidal effects of ROS^75^. We therefore asked if the antioxidant response that is driven by Nup214-dependent Nrf2 nuclear accumulation was the functional mechanism supporting *Anaplasma* persistence in *Ixodes* ticks. To address this, *nup214* expression was silenced through RNAi, and ISE6 cells were infected with *Anaplasma.* Cells were then supplemented with either vehicle control or the antioxidant N-acetyl cysteine (NAC) to see if the bactericidal phenotype could be rescued. We found that the addition of NAC restored *A. phagocytophilum’s* ability to infect and survive in tick cells when *nup214* is knocked down (Fig 5D). Altogether, our findings support a model where *Anaplasma* infection activates Nrf2, which drives the expression of Nup214. This leads to a positive feedback loop where Nup214 promotes nuclear accumulation of Nrf2, thereby enhancing antioxidant defenses that support *Anaplasma* persistence in ticks (Fig 6).

**Figure 6.**
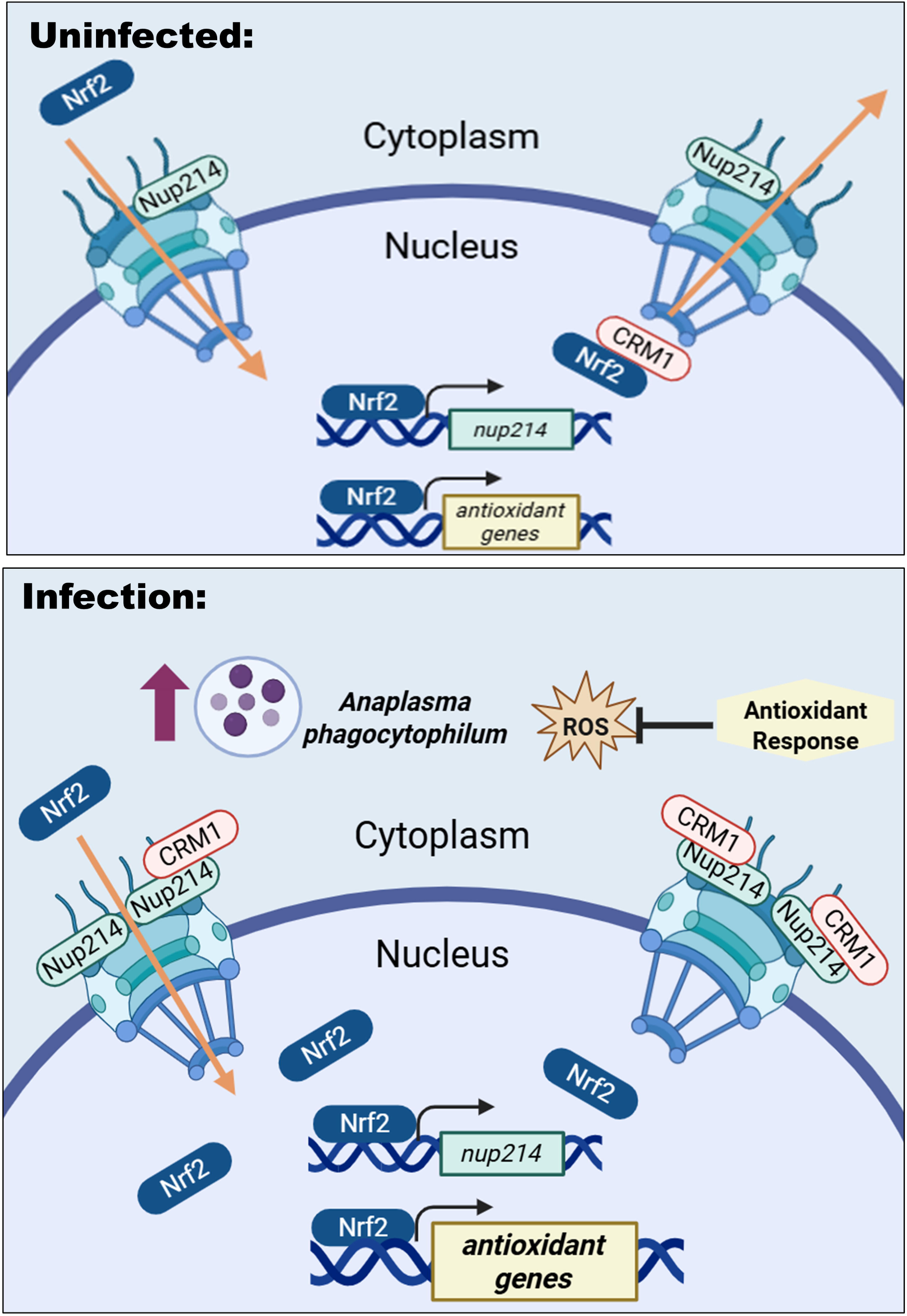
Nrf2-driven Nup214 expression amplifies the tick antioxidant response and supports *Anaplasma*. Infection activates the antioxidant response transcription factor Nrf2, which translocates to the nucleus and drives expression of *nup214*. Increased expression of Nup214 traps the nuclear export receptor CRM1 on the cytoplasmic side of the nucleus, which blocks export of Nrf2 and functionally amplifies the antioxidant response, benefitting *Anaplasma*.

## DISCUSSION

The ability of *Anaplasma* to persist in ticks can be attributed to both pathogen-mediated strategies and host-driven responses^7,13,76^. In ticks, initiation of the PERK pathway supports *Anaplasma* by activating Nrf2, which transcriptionally induces antioxidant defenses that reduce ROS^13^. Here, we show that Nrf2 also transcriptionally upregulates *nup214* in ticks. We demonstrate that increased Nup214 supports *Anaplasma* by trapping Nrf2 in the nucleus and amplifying the antioxidant response. Overall, this indicates a positive feedback loop, thereby creating a hospitable environment for *A. phagocytophilum*.

Ticks are obligate hematophagous arthropods that are reliant on host blood as a source of nutrients for growth and development. During digestion, large quantities of heme and iron are released, which promotes the production of ROS. To counter this threat, ticks have evolved robust antioxidant responses to protect themselves against oxidative damage. Whether *Anaplasma* is actively hijacking this system for its advantage or if it is passively benefiting from this host-driven response is not yet clear. However, targeting the NPC is a common strategy among pathogens to take advantage of host pathways and machinery for survival and replication^77–79^. Moreover, there are several examples of pathogens benefiting from Nup214, specifically. Adenoviruses and Herpesviruses use Nup214 as a docking site to import DNA to the nucleus^80,53,55^. The Rev protein from human immunodeficiency virus (HIV) is responsible for shuttling viral RNA out of the nucleus and uses Nup214 for CRM1-dependent export^54^. The obligate intracellular bacterial pathogen *Chlamydia trachomatis* secretes the effector protein CebN, which blocks STAT1 nuclear import by binding to nucleoporins, including Nup214. This functionally represses an interferon-γ-mediated immune response, thereby allowing the bacterium to infect and survive^56^. Our findings highlight another intracellular pathogen that benefits from Nup214, which may suggest that targeting the NPC is a common strategy among intracellular bacteria.

It is possible that *Anaplasma* could benefit from Nup214 in additional ways beyond regulating nuclear export. Most proteins enter the nucleus through associations with nuclear transport receptors that bind to nuclear localization signals (NLS) on the protein^81^. However, the FG domain of Nup214 can directly facilitate protein transport independently of nuclear transport receptors^61,82–84^. For example, proteins lacking classical NLSs such as SMAD2 (Mothers Against Decapentaplegic Homolog 2), ERK1/2 (Extracellular Signal-regulated kinase 1/2), and MAPK1 (Mitogen-activated protein kinase 1) all use Nup214 for nuclear import^82,84^. Many bacterial effectors that target the nucleus also lack a classical NLS. Instead, they contain either a non-classical NLS or they bind directly to nucleoporins for import^79^. There are several predicted, uncharacterized *A. phagocytophilum* effectors that are expressed during tick infection that contain either a classical or non-classical NLS^85,86^. It is therefore possible that Nup214 may be benefiting *Anaplasma* through an additional role by facilitating nuclear import of effectors.

To our knowledge, this is the first report that Nup214 expression impacts Nrf2 nuclear retention. However, it is well established that Nup214 regulates CRM1-dependent nuclear export^47,48,59,65,67^ and that other transcription factors are reliant on CRM1 to exit the nucleus^87^. Given this role, it is possible that Nup214 could lead to nuclear retention of additional infection-relevant transcription factors in *Ixodes,* thereby influencing *Anaplasma* infection dynamics.

Previously, we found that *Anaplasma* induces Nrf2 activation in ticks downstream from the UPR receptor, PERK^13^. In *Rosche et al.,* we knocked *PERK* expression down by RNAi in larvae and then fed them on either *Anaplasma* or *Borrelia-*infected mice. We found that, when *PERK* was knocked down, significantly less bacteria colonized and survived in the ticks during the early stages of infection compared to larvae that were treated with scrambled controls. After replete larvae molted, we found that there were still significantly less *Borrelia* in the resulting flat nymphs. However, *Anaplasma-*infected nymphs had bacterial numbers recover to levels that were comparable to the scrambled controls. In context of our current findings, increased Nrf2 nuclear retention through Nup214 could explain why *Anaplasma* numbers were able to rebound during the later stages of infection.

By using multiple methodologies, we uncovered a positive feedback loop linking Nrf2 and Nup214 that functionally amplifies the antioxidant response and influences infection outcomes. However, this is only one aspect of the Nrf2 transcriptional network in ticks. Our findings predict that there are many more genes targeted by Nrf2 that are unique when compared to the human network. Moreover, 28% of the genes putatively regulated by Nrf2 in *Ixodes* do not have human orthologs and remain uncharacterized (Supplementary Dataset 1). This highlights that there is still much to be studied about the *Ixodes* Nrf2 network that potentially extends beyond the antioxidant response and may be influencing vector competency for transmissible microbes.

## METHODS

### Cell cultures

The human embryonic kidney cell line, HEK 293T, was cultured at 33°C with 5% CO_2_ in Dulbecco’s modified Eagle medium (DMEM; Sigma, D6429) supplemented with 10% heat-inactivated FBS (fetal bovine serum) (Atlanta Biologicals, S115500) and 1x GlutaMAX (Gibco, 35050061).

*I. scapularis* embryonic ISE6 and IDE12 were cultured at 32°C with 1% CO_2_ in T-75 unvented tissue culture flasks (Fisher, 07-000-228). ISE6 and IDE12 cells were cultured in L15C300 media and L15C media, respectively, supplemented with 10% FBS (Sigma, F0926), 10% tryptose phosphate broth (BD, B260300), and 0.1% lipoprotein bovine cholesterol (MP Biomedicals, 219147680). Cell numbers and viability were measured with a trypan blue exclusion assay and a TC20^TM^ automated cell counter (Bio-Rad).

### Bacteria and animal models

*Escherichia coli* cultures were grown in lysogeny broth (LB) supplemented with ampicillin at 100 µg µl^-1^ with shaking between 230 and 250 rotations per minute (rpm) at 37°C overnight.

*A. phagocytophilum* strain HZ was cultured in Human Leukemia 60 (HL60) cells (American Type Culture Collection, CCL-240) in Roswell Park Memorial Institute 1640 (Cytiva, SH30027.LS) medium supplemented with 10% heat-inactivated FBS and 1x GlutaMAX. Cultures were maintained at 37°C with 5% CO_2_. HL60 cultures were kept between 1 x 10^5^ and 1 x 10^6^ cells mL^-1^.

*I. scapularis* larvae were acquired from Oklahoma State University (Stillwater, OK, USA). Ticks were maintained with 16:8 hour light:dark photoperiods and 95 to 100% relative humidity at 23°C.

Male C57BL/6 mice were obtained at ages 6 to 10 weeks old from colonies at Washington State University. Mice were infected intraperitoneally with 1 x 10^7^ host cell-free bacteria in 100 µl of phosphate buffered saline (PBS) (Intermountain Life Sciences, BSS-PBS) as previously described^13,88^. *A. phagocytophilum* burdens of each mouse were examined 6 days post-infection by collecting 25 to 50 µL of blood from the lateral saphenous vein, as previously described^13,22,88^. Bacterial burdens were quantified by quantitative real-time PCR (qRT-PCR) (*16s* relative to mouse β*-actin)*. All experiments with mice were conducted according to the guidelines and protocols approved by the American Association for Accreditation of Laboratory Animal Care (AAALAC) and by the Office of Campus Veterinarians at Washington State University (Animal Welfare Assurance A3485-01). The mice were housed in an AAALAC-accredited facility at Washington State University in Pullman, WA. All procedures were approved by the Washington State University Biosafety and Animal Care and Use Committees.

### RNAi silencing and pharmacological treatments

To synthesize silencing RNAs (siRNA) and scrambled RNAs (scRNA), the Silencer siRNA construction Kit (Invitrogen, AM1620) was used. ISE6 cells were seeded in a 24-well tissue culture plate at 1 x 10^6^ cells per well. 3 µg of siRNA or scRNA were transfected into tick cells with 2.5 µL of Lipofectamine 3000 (Invitrogen, L3000008) for 24 hours. Plates were spun at 450 x g for 1 hour as previously described^7^. 24 hours post-transfection, cells were infected with *A. phagocytophilum* at a multiplicity of infection (MOI) of 50 for 24 hours. The next day, cells were collected in TRIzol (Invitrogen, 1559026) for RNA isolation using the Direct-zol RNA Micropep Kit (Zymo, R2062). Complementary DNA (cDNA) was synthesized from 300 to 500 ng total RNA using the Verso cDNA Synthesis Kit (Thermo Fisher Scientific, AB1453B). Silencing efficiency, bacterial burden, and gene expression were assessed by qRT-PCR with iTaq universal SYBR Green Supermix (Bio-Rad 1725125) (Table S1). Cycling conditions followed manufacturer’s recommendations.

For NAC treatment, cells were seeded at 1 x 10^6^ cells per well in a 24-well plate and transfected with siRNA for *nup214* or the scrambled control as described above. Cells were infected with *A. phagocytophilum* (MOI 50) for 24 hours alone or in the presence of 50 mM NAC (Sigma, A7250). Cells were collected in TRIzol for RNA extraction as described above. Bacterial burden and silencing efficiency were assessed by qRT-PCR as described above.

For Leptomycin B (LMB) treatment, cells were seeded at 2.5 x 10^5^ cells per well in a 96-well plate and infected with *A. phagocytophilum* (MOI 50). At 20 hours, cells were treated with 20 nM of LMB (Sigma, L2913) or equal volume of ethanol. Cells were collected at 3 hours post treatment for cell viability assessment or for RNA extraction as described above. qRT-PCR was used to assess bacterial burden. All data are expressed as means ± standard error of the mean (SEM).

### Protein structure predictions and alignments

*Ixodes* Nup214 and *Ixodes* CRM1 were identified using NCBI Protein Basic Local Alignment Tool (BLAST) and querying the tick genome with human protein sequences (Nup214: NP_005076.3, CRM1: NP_001397728.1) as done previously for *Ixodes* Nrf2^13^. Jalview was used to visualize all protein alignments^89^. Physiochemical property conservation between amino acids was indicated by shading. AlphaFold3 was used to model *Ixodes* Nrf2 and *Ixodes* MafG binding to the *Ixodes nup214* promoter^58^ and visualized in UCSF ChimeraX^90^. Input sequences used for AlphaFold3 are found in Supplementary Dataset 2.

### Predicting Nrf2 binding sites in the tick genome

All binding site analyses were conducted as previously described using ArthroQuest^7^. Briefly, putative promoter regions of current genomes available on ArthroQuest were scanned for the following Nrf2 binding motif: [A/G]TGA[C/G/T][T/A]n[A/G]GC, where “n” is any nucleotide^32^. The resulting data included predicted promoter sites with the binding site detected and corresponding gene annotations. Using a script previously described in Vosbigian *et al*. (2025)^7^, human (GCF_000001405.40) and *Drosophila* (GCF_000001215.4) orthologs of the putative Nrf2-regulated genes were identified in R. All ortholog inquiries were conducted using R version 4.2.2 and RStudio 2022.07.2.576^91,92^. Human and *Drosophila* orthologs were then queried in Gene Ontology and Reactome databases^93,94^.

### Plasmid construction

The *nup214* promoter sequence was amplified via PCR from ISE6 DNA with primers listed in Table S1. The mutated Nrf2 promoter was synthesized by GenScript and was changed from “GTGATTAAGC” to “TGCTAGCTTA**”**. The promoter sequences were cloned into a pTE luciferase reporter (Signosis, LR-2200) using *BgIII. Ixodes* Nrf2 was synthesized by GenScript and was codon optimized for mammalian and *E. coli* expression. *Ixodes Nrf2* was cloned into pCMV-HA (New MCS) (gift from Christopher A. Walsh; Addgene plasmid number 32530) using *EcoRI* and *EcoRV*. The C-terminal truncation of Nrf2 (2101-3003) was amplified from the Nrf2 plasmid using primers (Table S1) and cloned back into pCMV-HA. *Ixodes* CRM1 (AA 1603-3310) was amplified from ISE6 cDNA with primers (Table S1) via PCR and cloned into pCMV/hygro-Negative control vector (SinoBiological; CV005) using *HindIII* and *KpnI*. All constructs were confirmed by sequencing through Plasmidsaurus.

### HEK 293T cell transfection

In 6 well plates, HEK 293T cells were seeded at 1 x 10^6^. The following day, cells were transfected with pCMV-Nrf2-HA and pCMV-CRM1-FLAG plasmid DNA using 10 µL Lipofectamine 3000 and 10 µL P3000 reagent (Fisher Scientific, L30000015), in Opti-MEM I reduced-serum medium (Gibco, 31985062). After 5 hours, the medium was replaced with complete DMEM and incubated at 33°C, 5% CO_2_ for 24 hours. Cells were collected in RIPA (radioimmunoprecipitation assay; Fisher Scientific; P189901) supplemented with 1x protease and phosphatase inhibitor cocktail (Thermo Scientific, 78440)

### Polyacrylamide gel electrophoresis and western blotting

Bicinchoninic acid (BCA) protein assays (Pierce, 23225) determined protein concentrations. For each sample, 20 µg of protein was separated at 100 V for approximately 1 hour and 30 minutes using a 4 to 15% Mini-PROTEAN TGX (Tris-Glycine eXtended) precast gel (Bio-Rad, 4568084). Proteins were transferred to a polyvinylidene difluoride membrane. Membranes were blocked in 5% milk in 1 x phosphate-buffered saline containing 0.1% Tween 20 (PBS-T) for 0.5 to 2 hours at room temperature. Primary antibodies were diluted in PBS-T with either 5% milk or 3% Bovine Serum Albumin and incubated overnight at 4°C. The primary antibodies included anti-HA (Invitrogen, 26153, 1:1000), anti-FLAG-HRP (Sigma, A8512; 1:2000), anti-Nrf2 (OriGene, OTI8A10; 1:1000), and anti-Actin (Sigma, A2105; 1:1000).

### Coimmunoprecipitation assays

Expression of *Ixodes* Nrf2-HA (aa 701-1000) and CRM1-FLAG (aa 1603-3310) was validated with anti-HA (Pierce; 26183; 1:1,000) and anti-FLAG-HRP (Sigma; A8592; 1:500). Cross-linked agarose beads (anti-FLAG M2 [Sigma; A2220] and anti-HA [Pierce; 26181]) were washed three times with wash buffer (25mM Tris-HCl, 150 mM NaCl, 1% NP-40, 1mM EDTA; pH 7.5) and incubated with wash buffer at 4°C for 1 hour. 0.4 mg of cell lysate was brought to a final volume of 1 mL in wash buffer containing protease and phosphatase inhibitors, combined with 80 µL (packed volume) of cross-linked agarose beads and incubated overnight at 4°C with rotation. Beads were washed 3 times with wash buffer, and protein was eluted by boiling in 50 µL of 4x Laemmli buffer for 5 minutes. Protein interactions were evaluated by immunoblotting as described above.

### Nuclear extractions

Nuclear extracts of ISE6 tick cells were obtained using the NE-PER Nuclear and Cytoplasmic Extraction Kit (Thermo Scientific, 78835). 1 x 10^6^ tick cells per well were seeded in 12-well plates. Tick cells were either infected with *Anaplasma phagocytophilum*, silenced for *nup214* with RNAi, or treated with LMB according to protocols described above. Once nuclei were collected, 20 µg of nuclear extracts were then analyzed by immunoblotting as described above.

### Luciferase reporter assays

To assess *nup214* regulation by Nrf2, 1 x 10^4^ HEK 293T cells were seeded in clear-bottom, white-walled 96-well plates (Greiner Bio-One, 655098). The next day, cells were transfected with 0.05 µg of the *nup214* reporter plasmid or mutated *nup214* reporter plasmid in combination with a plasmid expressing *Ixodes* Nrf2 using 0.5 µL of Lipofectamine 3000 in Opti-MEM I reduced serum medium (Gibco, 31985062). After 5 hours, the medium was replaced with complete DMEM and incubated at 33°C, 5% CO_2_ for 24 hours. Luminescence was measured by adding 5 mg mL^-1^ of D-Luciferin potassium salt (Promega, E1500) to each well and quantifying with a plate reader. Data are represented as relative luciferase units (RLU) normalized to control condition containing only the reporter plasmid without *Ixodes* Nrf2 expression plasmid ± SEM.

### Gene expression analysis of whole ticks

*I.* scapularis larvae were fed until repletion on either uninfected or *A. phagocytophilum*-infected mice. Larvae were then collected and either immediately processed or maintained in an incubator for 7 days post-repletion. Prior to adding TRIzol, individual ticks were flash frozen in liquid nitrogen and pulverized mechanically. RNA was isolated and cDNA was synthesized as described above. Using the primers listed in Table S1, gene expression was measured by qRT-PCR. All samples were normalized to uninfected controls. Data are expressed as means ± SEM.

### RNAi silencing and analysis of whole ticks

As previously described, *I. scapularis* larvae were silenced with RNAi^7,13,22^. Around 100 larvae were added to a 1.5 mL tube with 40 µL of siRNA or scrambled controls and incubated at 15 °C overnight. Larve were dried with filter paper and allowed to recover overnight before being placed on mice. Over 3 to 5 days post-tick placement, replete larvae were collected and frozen. Larvae were weighed in groups of 3 to assess feeding efficiency. RNA was isolated from each tick as described above. qRT-PCR was performed with a standard curve to generate absolute numbers of target sequences. Standard curves were generated with a plasmid containing either *A. phagocytophilum 16s*, *Ixodes* β*-actin*, or *Ixodes nup214* (Table S1).

### Reactive oxygen species and antioxidant assays

For measuring ROS, ISE6 cells were seeded at 1.68 x 10^5^ per well in black walled-clear bottom 96-well plates (Thermo Scientific, 165305). Cells were transfected with 2.25 μg of siRNA for *nup214* or the scrambled control as described above. 24 hours later, all wells were treated with DCF-DA, the fluorescent detection probe (10 µM; Sigma, D2926) for 1 hour in Ringer buffer (155 mM NaCl, 5 mM KCl, 1 mM MgCl_2_ · 6H_2_O, 2 mM NaH_2_PO_4_ · H_2_O, 10 mM HEPES, and 10 mM glucose). After the buffer was removed, the cells were washed with PBS at room temperature and *A. phagocytophilum* (MOI 50) was added. Fluorescence was measured at 504 nm/ 529 nm at 24 hours post-infection. Data were graphed as fold change of relative fluorescence units (RFU) normalized to the uninfected, scrambled control ± SEM.

For measuring the activity of the antioxidant, Glutathione-S-Transferase (GST), we adapted a GST activity assay protocol for tick cells^95–97^. After washing in PBS, cells were resuspended in sodium phosphate buffer (200 μL, pH 7.2, 50 mM) and sonicated (4 x 15 seconds, amplitude: 30). 10 μL of each sample was added to sodium phosphate buffer in white-walled clear clear-bottom 96-well plates (Greiner Bio-One, 655098). 1-chloro-2,4-dinitrobenzene (CDBN; 30 mM; 10 μL) and reduced glutathione (GSH; 0.1M; 2μL) were added, followed by immediately placing in the plate reader. Absorbance was measured at 340 nm every 2 minutes for 10 minutes total. GST activity was calculated according to the protocol for the GST Assay Kit (Sigma; CS0410). GST activity was normalized to the uninfected, scrambled control.

### Statistical analysis

*In vitro* experiments included at least 3 to 5 replicates. *In vivo* experiments included at least 10 to 20 ticks. Non-parametric Mann-Whitney test or an unpaired Student’s *t* test expressed as means ± SEM were used to assess data. Graphs were created in GraphPad Prism. Statistical significance was determined by a *P* value of <0.05.

## ACKNOWLEDGEMENTS

We are grateful to Ulrike Munderloh (University of Minnesota) for providing the ISE6 tick cell line and Oklahoma State University for *Ixodes scapularis* ticks. The Addgene plasmid 32530 was a gift from Christopher A. Walsh. Schematics in Fig 1 and 6 were created with BioRender.

## Author Contributions

K.A.V. and D.K.S. designed the study. K.A.V., C.A.O., B.P.S., K.L.R., S.J.W., and E.R.Z. performed experiments. K.A.V. and D.K.S. analyzed data. All authors provided intellectual input into the study. K.A.V. and D.K.S. wrote the manuscript. All authors contributed to editing.

## FIGURE LEGENDS

**<u>Supplementary Figure 1.</u> Nrf2 antibody validation.** (A) Mammalian Nrf2 antibodies tested against *Ixodes* Nrf2 (predicted size 107 kDa, but migrates at ∼150 kDa). (B) Immunoblot against HEK 293T cells transfected with HA-tagged pCMV *I. scapularis* Nrf2.

**<u>Supplementary Figure 2.</u> *gst theta-1* expression during Nrf2 knockdown.** *gst theta-1* expression quantified from *A. phagocytophilum* infected IDE12 tick cells treated with siRNA targeting *nrf2* or a scrambled control. *, *P* < 0.05 (Student’s *t* test).

**<u>Supplementary Figure 3.</u> Associated pathways with *Ixodes* genes putatively regulated by Nrf2.** Orthologs of predicted Nrf2-regulated genes that are associated with Nrf2-regulated pathways in Reactome (teal) and Gene Ontology (GO, blue) databases.

**<u>Supplementary Figure 4.</u> *I. scapularis* MafG sequence alignment and structural predictions.** (A) Human and *I. scapularis* MafG amino acid sequences were aligned and visualized via Jalview. Conservation by percent identity is indicated by shading. (B) *I. scapularis* MafG predicted by AlphaFold3. Color of each residue represents the model confidence score, pLDDT. Blue to orange indicates high to low confidence, respectively. (C) *I. scapularis* MafG (purple) aligned with human MafG structure (orange). (D) AlphaFold3 predicted DNA complex of *I. scapularis* Nrf2, *I. scapularis* MafG, and *I. scapularis nup214* promoter. Color of each residue represents pLDDT.

**<u>Supplementary Figure 5.</u> Luciferase reporter assay validation**. HEK 293T cells were co-transfected with plasmids constitutively expressing *I. scapularis* Nrf2 and a commercially available luciferase reporter plasmid containing the 8 consecutive Nrf2 binding sites. *, *P* < 0.05 (Student’s *t* test).

**<u>Supplementary Figure 6.</u> Nup214 sequence alignment and structure predictions with the human ortholog.** (A) Human and *I. scapularis* Nup214 amino acid sequences were aligned and visualized via Jalview. Conservation by percent identity is indicated by blue shading, green shading indicates FG repeats and orange boxes indicate CRM1 binding sites (*H. sapiens Nup214* 1916-1951, 1980-1988, 2009-2027). (B) *I. scapularis* beta-propeller domain of Nup214 predicted by AlphaFold3. Color of each residue represents the model confidence score, pLDDT. Blue to orange indicates high to low confidence, respectively. (C) Alignment of the Nup214 beta-propeller domain structure of human Nup214 (teal) and tick Nup214 (magenta).

**<u>Supplementary Figure 7.</u> CRM1 binding sites, sequence, and structure prediction alignment.** (A) *I. scapularis* Nrf2 sequence with CRM1 binding sites highlighted in purple. CRM1 truncation highlighted by magenta box. (B) Human and *I. scapularis* CRM1 amino acid sequences were aligned and visualized via Jalview. Shading indicates conservation by percent identity. Yellow indicates NES binding loop, teal box indicates the CRM1 truncation. (C) *I. scapularis* CRM1 predicted by AlphaFold3. Color of each residue represents the model confidence score, pLDDT. Blue to orange indicates high to low confidence, respectively. (D) Alignment of *I. scapularis* CRM1 (orange) with human CRM1 (blue).

**<u>Supplementary Figure 8.</u> AlphaFold3 model of *I. scapularis* Nrf2-CRM1 interaction.** (A) *I. scapularis* Nrf2-CRM1 interaction. Color of each residue represents the model confidence score, pLDDT. Blue to orange indicates high to low confidence, respectively. (B) *I. scapularis* Nrf2 (yellow) interactions with *I. scapularis* CRM1 (magenta). CRM1 binding groove highlighted in teal. Predicted NES sites in the bZIP domain of Nrf2 are highlighted in orange.

**<u>Supplementary Figure 9.</u> Tick cell viability with Leptomycin B.** Percent viability of ISE6 cells treated with 20 nM LMB for 3 hours. NS, not significant.

