## Supplemental Table 1 for "An Nrf2-Nup214 positive feedback loop sustains the antioxidant response and promotes microbial infection in ticks"

**Supplementary Table 1.** Oligonucleotide primers used in this study

| **Name** | **Target gene** | **Primer Sequences** |
| --- | --- | --- |
| Mus musculus β-Actin (qRT-PCR) | XM_030254057.1 | F 5'-ACGCAGAGGGAAATCGTGCGTGAC-3'  R 5'-ACGCGGGAGGAAGAGGATGCGGCAGTG-3' |
| Anaplasma phagocytophilum 16S (qRT-PCR) | NC_007797 | F 5'-CCCTAAGGCCTTCCTCACTC-3'  R 5'-CAGCCACACTGGAACTGAGA-3' |
| Anaplasma phagocytophilum 16S_full | NC_007797 | F 5'-TCCTGGCTCAGAACGAACG-3'  R 5'-GTCACTGACCCAACCTTAAATGG-3' |
| Ixodes scapularis Actin (qRT-PCR) | XM_029977298.1 | F 5'-GCCGGGACCTTACAGACTATC-3'  R 5'-CACGGACAATTTCACGCTCG-3' |
| Ixodes scapularis Nup214  (qRT-PCR) | XM_042292365.1 | F 5’-CTCGCAGCATCGTCTAGGTT-3’  R 5’-TCCAGCCGGTAGATCTTCAC-3’ |
| Ixodes scapularis Nrf2  (qRT-PCR) | XM_042293400.1 | F 5’-GTCTTCGACTTCCGGTTTGA-3’  R 5’-GTAGGCACTTCGGTGCTCTC-3’ |
| Ixodes scapularis Keap1  (qRT-PCR) | XM_029970525.4 | F 5’-TGAAGGAAACGGAAGAGTACC-3’  R 5’-CCTTCCGGTGGTACGTCC-3’ |
| Ixodes scapularis Crm1  (qRT-PCR) | XM_029966788.3 | F 5’- AGCAAGACCAATGAGAGCCT-3'  R 5’- GAAGTCAAACACCTCCTCGC-3' |
| Ixodes scapularis Nup214 siRNA_771 | XM_042292365.1 | F 5’-AACCAAACAACGGACTCCCAACCTGTCTC-3’  R 5’-AATTGGGAGTCCGTTGTTTGGCCTGTCTC-3’ |
| Ixodes scapularis Nup214 scRNA | N/A | F 5’-AAGACCAACAACCCCGTAACACCTGTCTC-3’  R 5’-AATGTTACGGGGTTGTTGGTCCCTGTCTC-3’ |
| Ixodes scapularis Nrf2 siRNA_1058 | XM_042293400.1 | F 5’-AACCTTCATGCATGGATCCTTCCTGTCTC-3’  R 5’-AAAAGGATCCATGCATGAAGGCCTGTCTC-3’ |
| Ixodes scapularis Nrf2 scRNA | N/A | F 5’-AAGCTTACCGAGTCTCCTATTCCTGTCTC-3’  R 5’-AAAATAGGAGACTCGGTAAGCCCTGTCTC-3’ |
| Ixodes scapularis Keap1 siRNA_95 | XM_029970525.4 | F 5’-AAGCATAGACACCTTTCCTAACCTGTCTC-3’  R 5’-AATTAGGAAAGGTGTCTATGCCCTGTCTC-3’ |
| Ixodes scapularis Keap1 scRNA | N/A | F 5’-AAGACTTATCCCACGTTAACACCTGTCTC-3’  R 5’-AATGTTAACGTGGGATAAGTCCCTGTCTC-3’ |
| Ixodes scapularis Crm1 siRNA_1065 | XM_029966788.3 | F 5’- AAGCAGCGAATTCTCCCACATCCTGTCTC-3'  R 5’- AAATGTGGGAGAATTCGCTGCCCTGTCTC-3' |
| Ixodes scapularis Crm1 scRNA | N/A | F 5’-AAGAACGTCTCTCCGAACCTACCTGTCTC-3'  R 5’-AATAGGTTCGGAGAGACGTTCCCTGTCTC-3' |
| pCMV-Nrf2-HA | XM_042293400.1 | F 5’-AAAAGAATTCATGATGAACTACAAAAAGTGCAGCG-3’  R 5’-AATTGATATCTTACTCATCTTCCTCCCATTTAATA-3’ |
| PCMV-Nrf2 (AA 2101-3003)-HA | XM_042293400.1 | F 5’ -AAAAGAATTCCTGCCGAGATTCCTGGGTAG-3'  R 5’-AATTGATATCTTACTCATCTTCCTCCCATTTAATA-3’ |
| pCMV-CRM1 (AA 1603-3310) -FLAG | XM_029966788 | F 5’- AACCAAGCTTATGCACGAGGAGGACGAGAAGC -3’  R 5’-ACTTGGTACCGTCCTGCATCTCTTCTGGGATCT-3’ |
| Pte-Nup214 Promoter-Luciferase | Predicted promoter region found in Table S2 | F 5’-ACGCAGATCTAATCCAGCGTGTCGGCAGC-3’  R 5’-ACGCAGATCTTGGCTTTAAGGTGCGAGCGAA-3’ |
| Ixodes scapularis GST-1 Theta (qRT-PCR) | XM_040208699.3 | F 5’-GCCTTCCGCTCACAATTAAG-3’  R 5’-ACAGTGTGGGCTGGGTTAAG-3’ |
