## Supplementary figures and images for "An Nrf2-Nup214 positive feedback loop sustains the antioxidant response and promotes microbial infection in ticks"

### Supplemental Figure 1

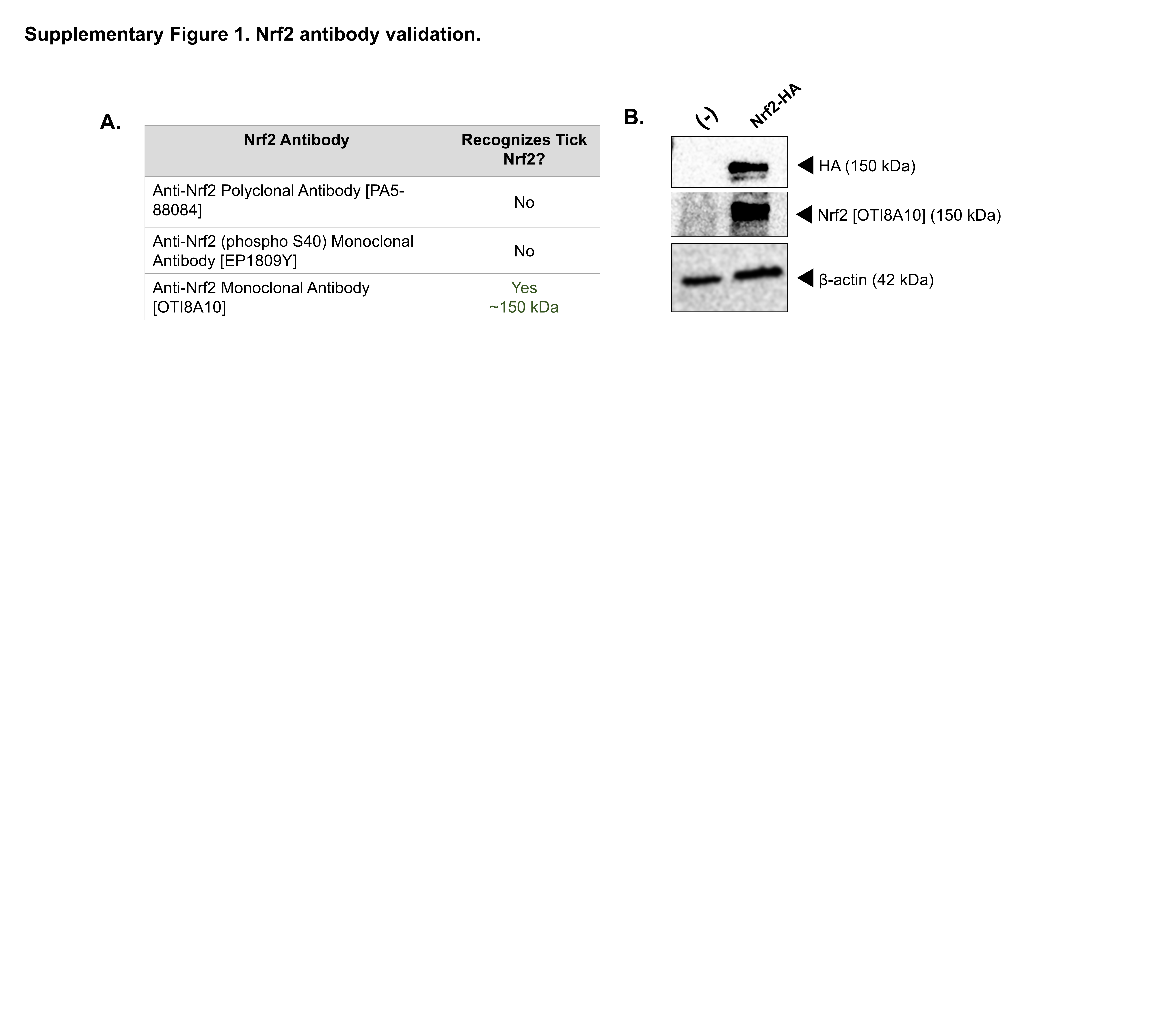

### Supplemental Figure 2

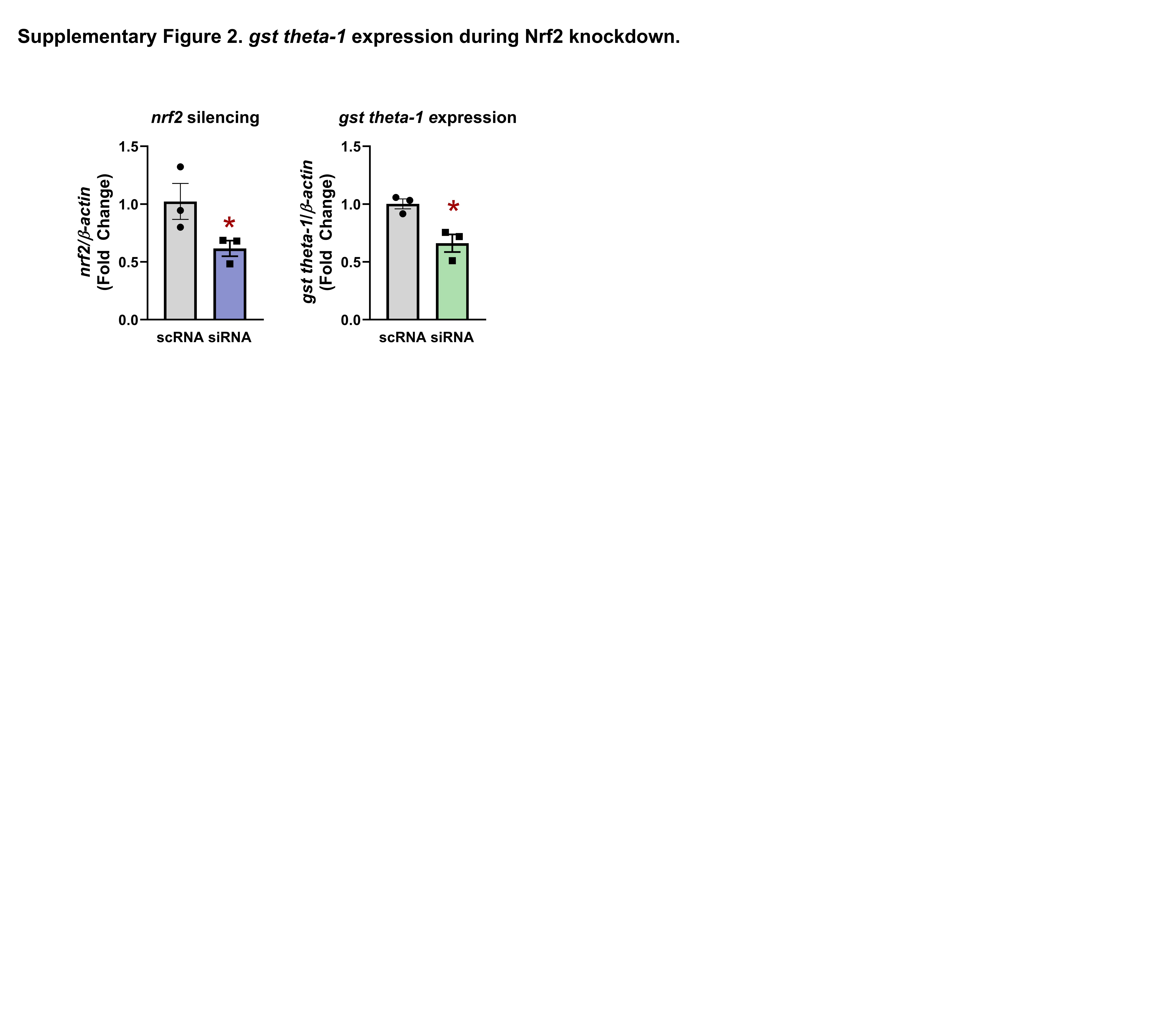

### Supplemental Figure 3

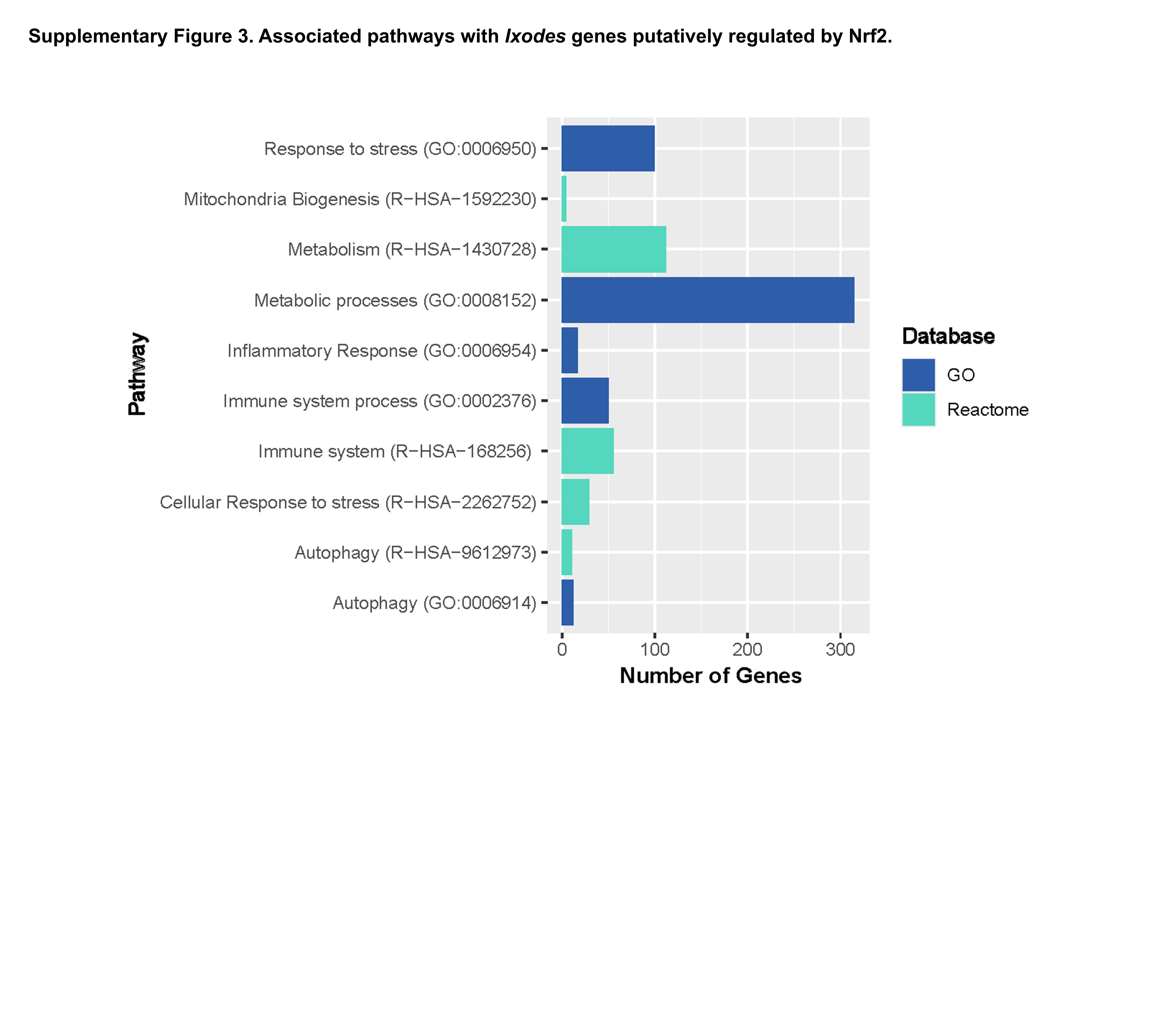

### Supplemental Figure 4

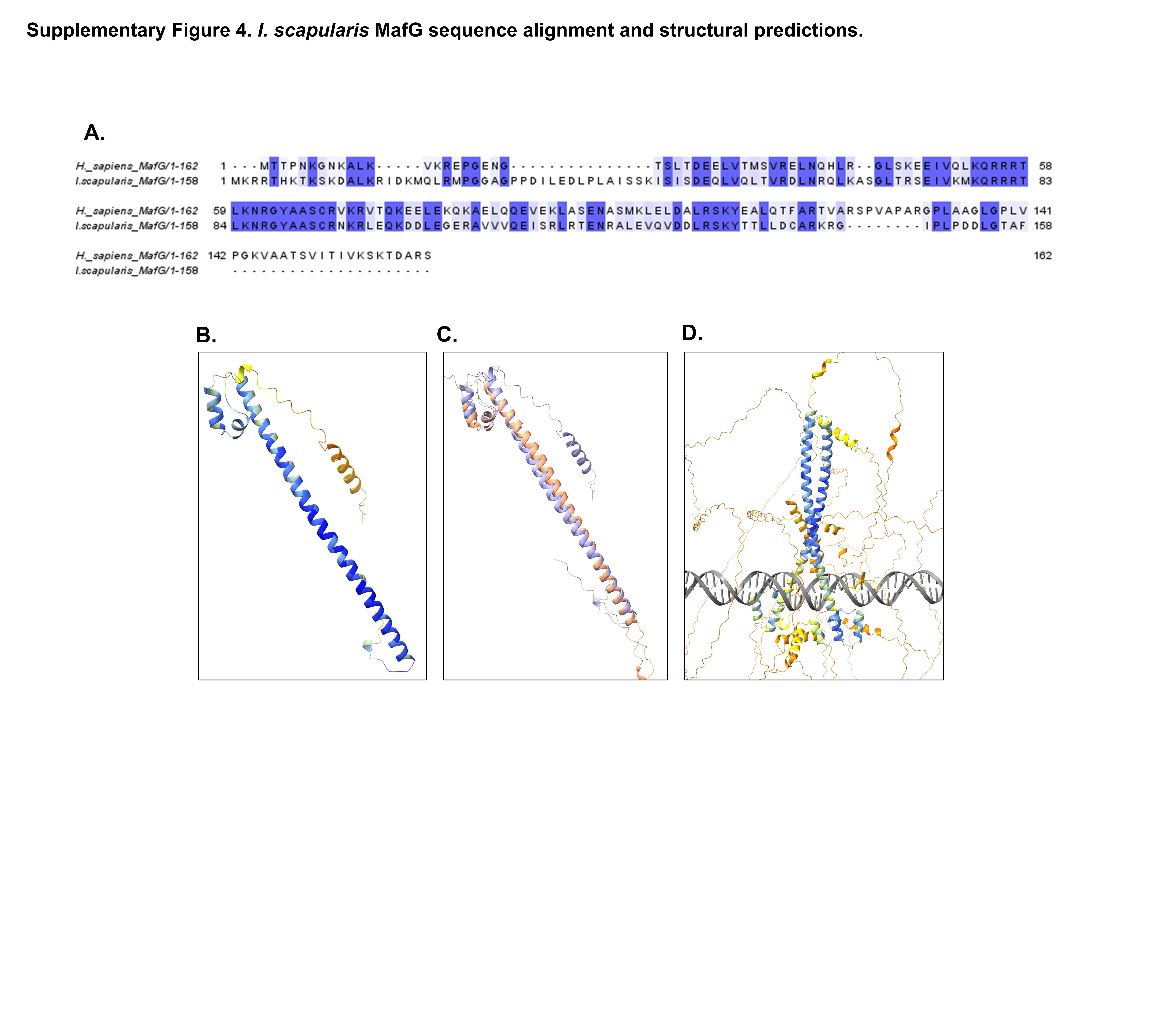

### Supplemental Figure 5

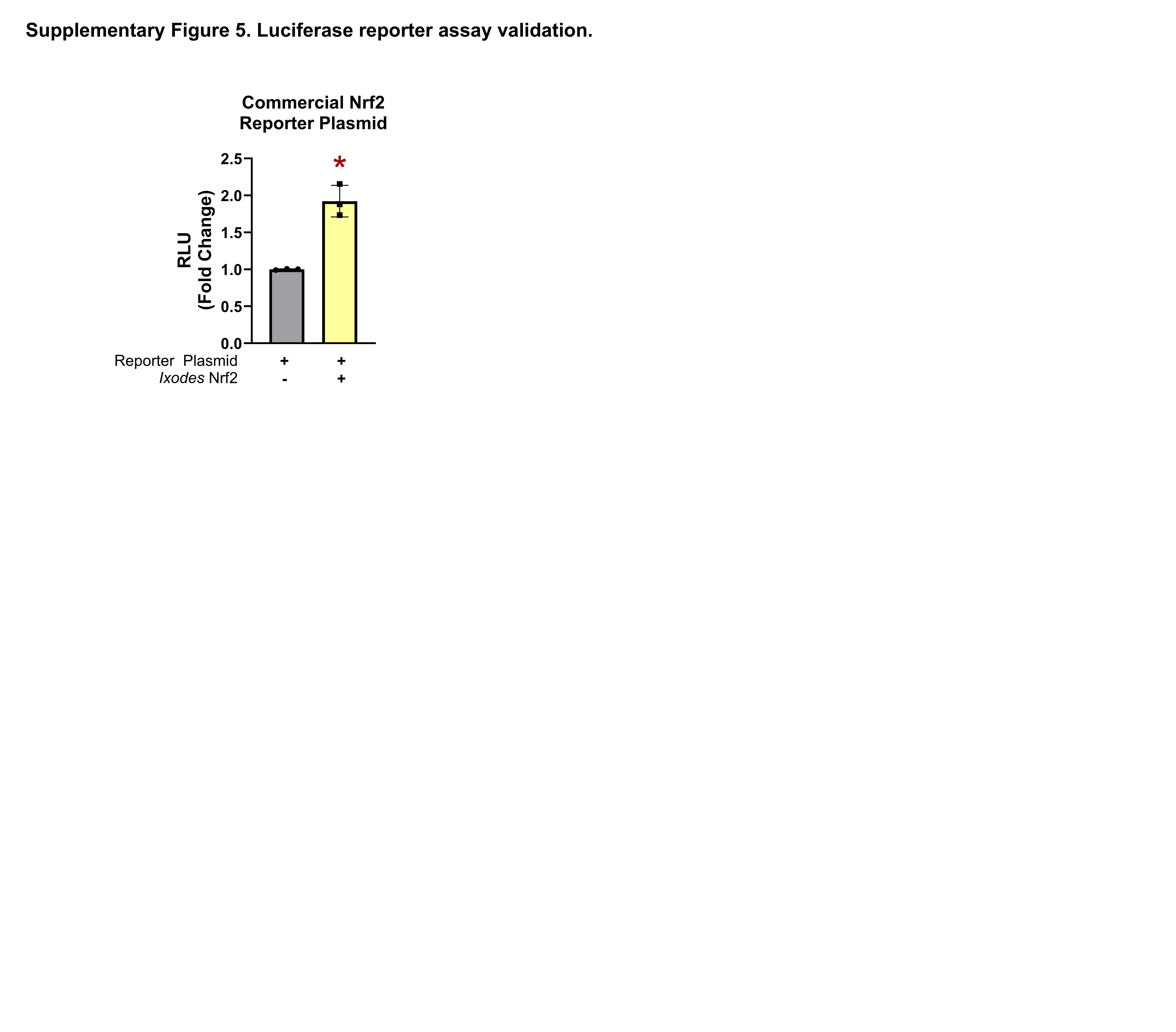

### Supplemental Figure 6

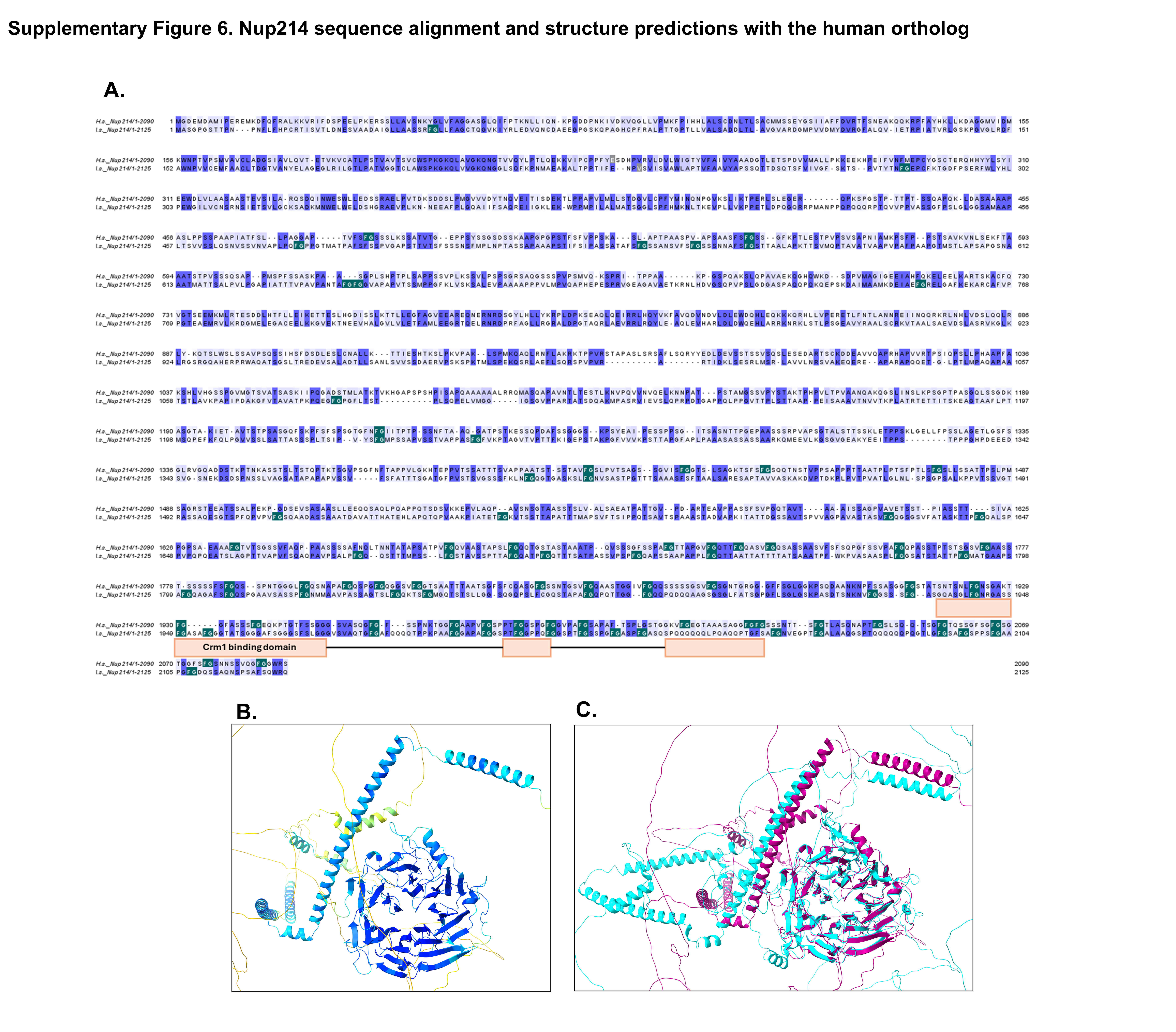

### Supplemental Figure 7

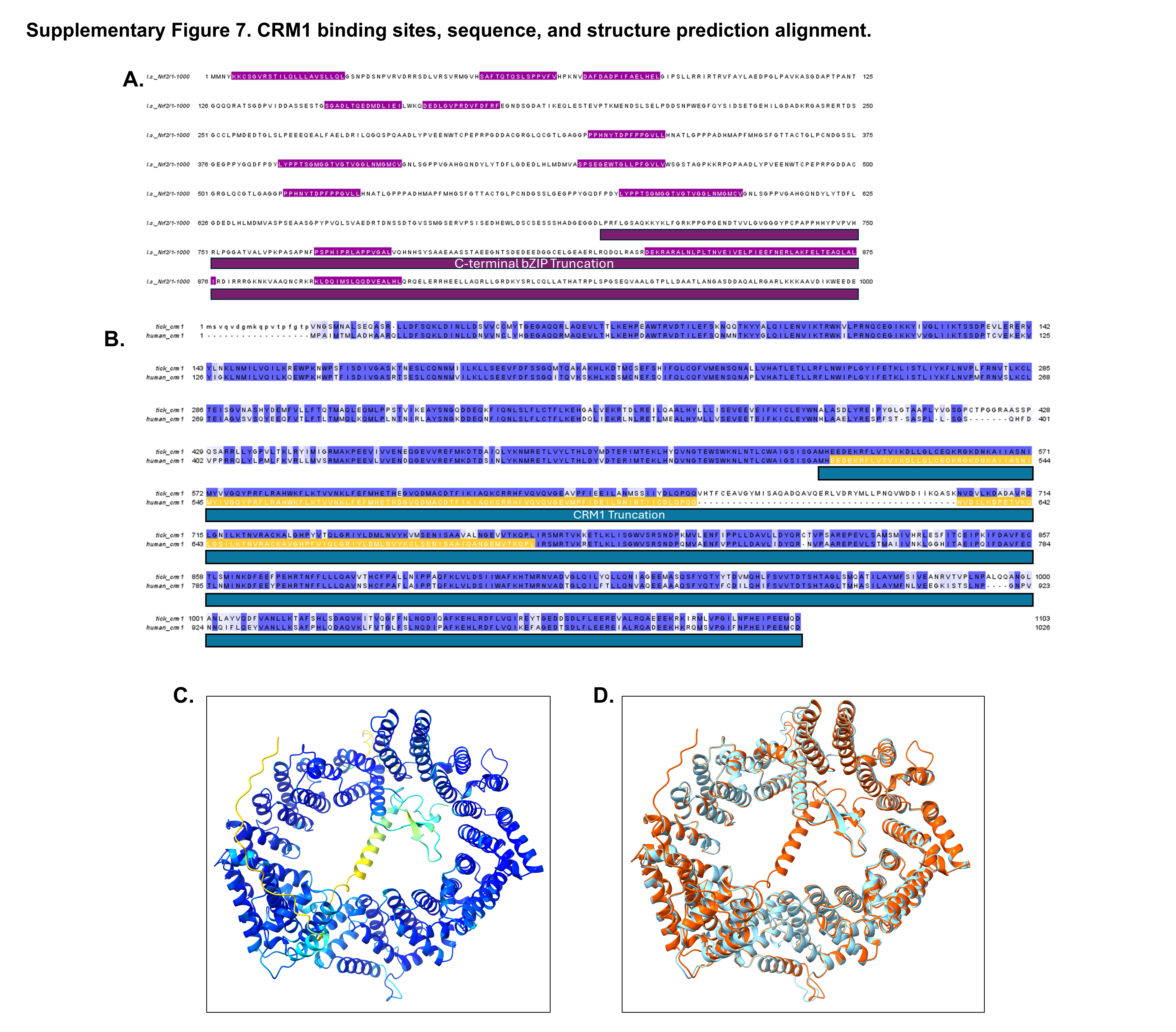

### Supplemental Figure 8

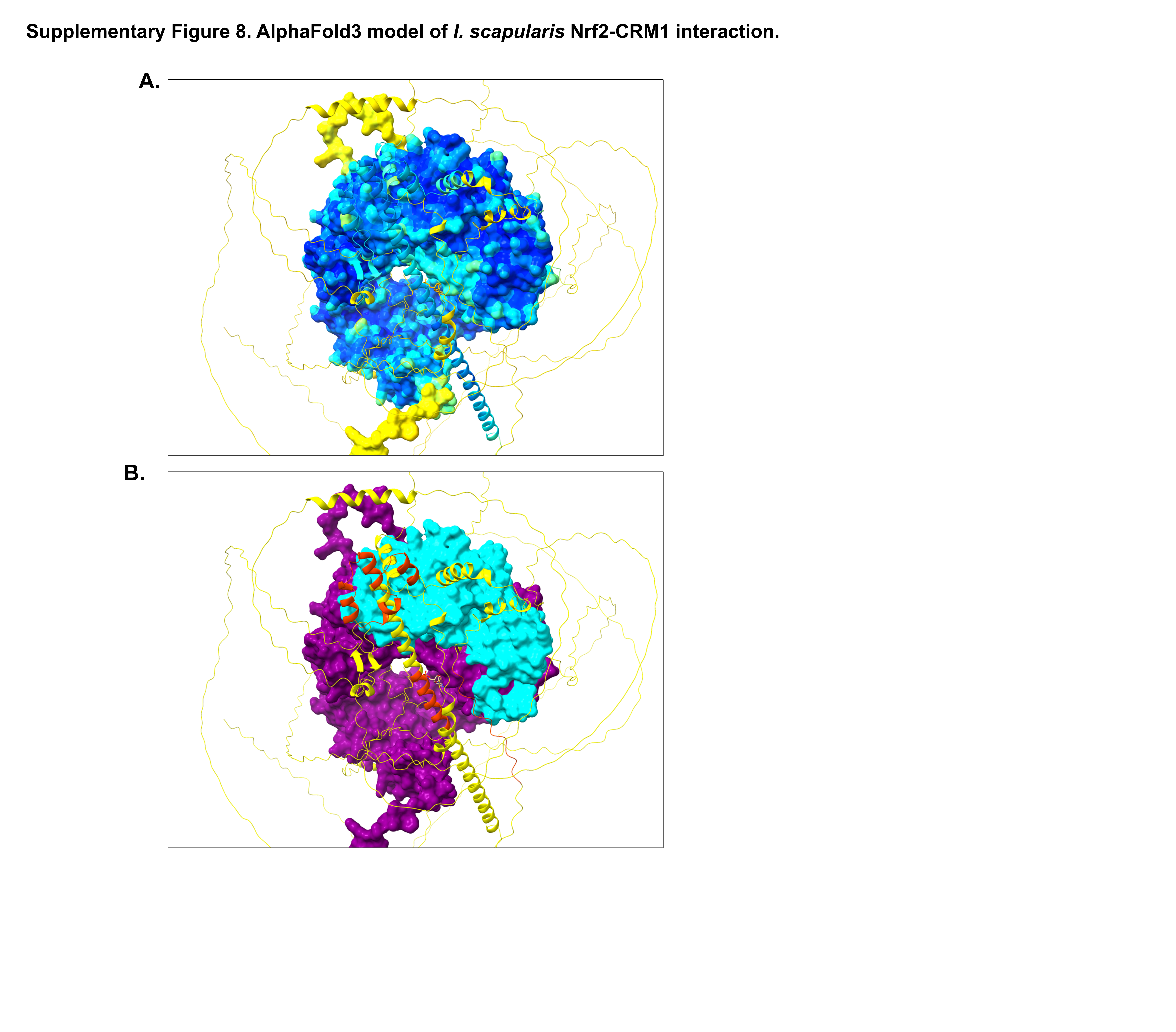

### Supplemental Figure 9

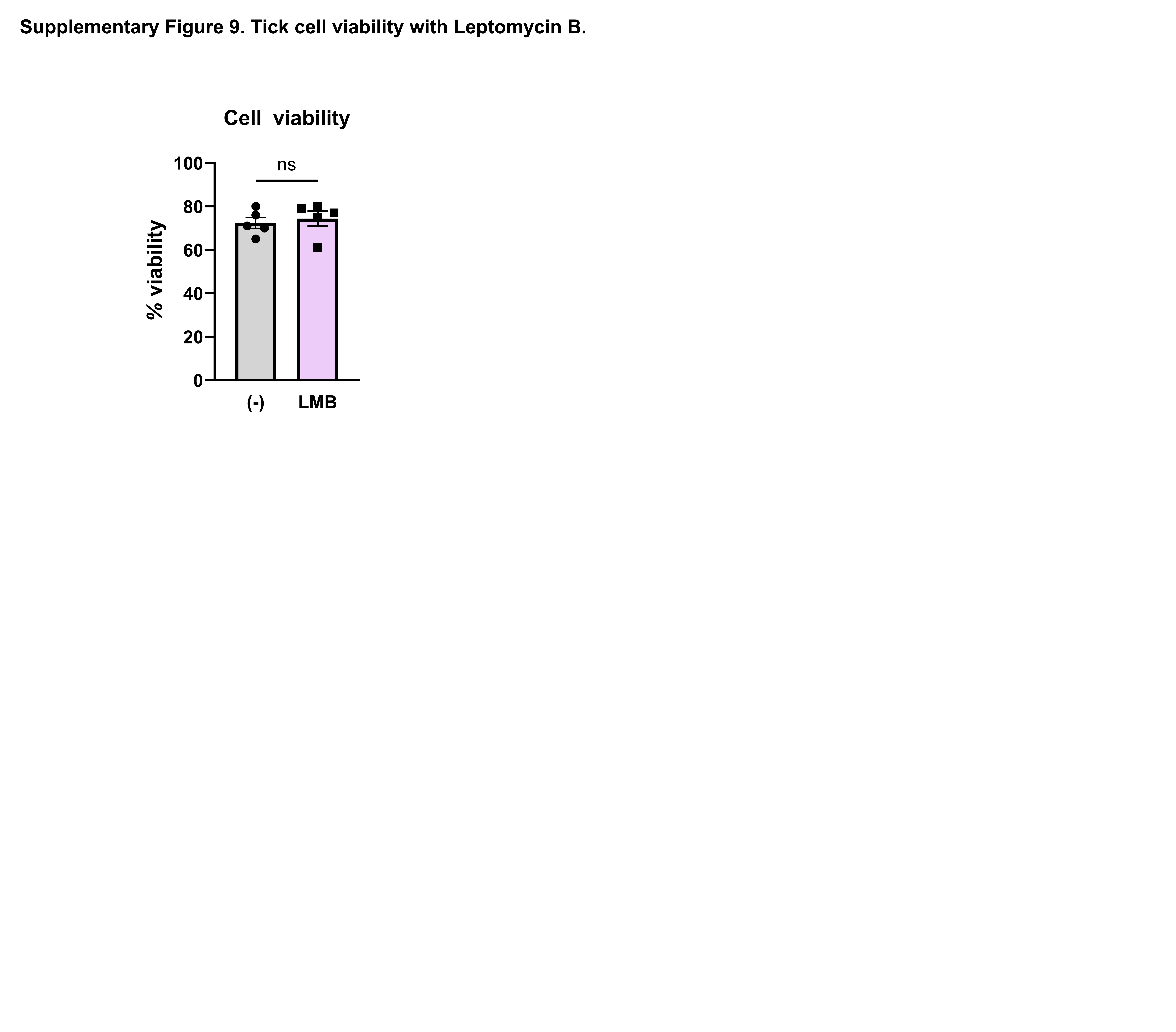
